# Distributed coding of stimulus magnitude in rodent prefrontal cortex

**DOI:** 10.1101/2020.04.02.021287

**Authors:** Josephine Henke, David Bunk, Dina von Werder, Stefan Häusler, Virginia L Flanagin, Kay Thurley

## Abstract

As we interact with the external world, we judge magnitudes from sensory information. The estimation of magnitudes has been characterized in primates, yet it is largely unexplored in non-primate species. Here, we show that gerbils that solve a time-interval reproduction task display primate-like magnitude estimation characteristics, most prominently a systematic overestimation of small stimuli and an underestimation of large stimuli, often referred to as regression effect. We investigated the underlying neural mechanisms by recording from medial prefrontal cortex and show that the majority of neurons respond either during the measurement or the reproduction of a time-interval. Cells that are active during both phases display distinct response patterns. We categorize the neural responses into multiple types and demonstrate that only populations with mixed responses can encode the bias of the regression effect. These results reveal the organizing neural principles of magnitude estimation a process important to higher cognitive functions.

## Introduction

Animals including humans estimate the magnitude of physical stimuli, integrate path length and keep track of duration to gather behaviorally relevant information from their environment. Although such estimates may ultimately be used for binary actions, like discriminating items or events and making decisions, the estimation itself is done on a continuum of values. Behavioral analyses over the past century established specific biases in magnitude estimation (e.g. reviewed in Petzschner, Glasauer, and Stephan, 2015) such as the *regression effect*, i.e. the overestimation of small and the underestimation of large stimuli across a range of values (also known as regression to the mean, central tendency, or Vierordt’s law). Recently, this bias regained attention as it may be the result of an error minimization strategy (Cicchini et al., 2012; Jazayeri and Shadlen, 2010; Petzschner and Glasauer, 2011).

Despite a long history of behavioral research on magnitude estimation, its neural basis is not well understood. It is an ongoing debate whether a dedicated or distributed magnitude system exists in the brain (for review see Bueti and Walsh, 2009; Kadosh, Lammertyn, and Izard, 2008; Van Opstal and Verguts, 2013). Human studies identified frontal, parietal and striatal brain regions that are active during magnitude estimation. Recent studies in non-human primates found that neural population dynamics in frontal and parietal cortices subserve magnitude estimation behavior (Jazayeri and Shadlen, 2015; Sohn et al., 2019). These studies investigated the estimation of time intervals lasting hundreds of milliseconds. It is still unresolved how their findings translate to durations of several seconds, i.e. to time scales that are relevant for more complex and ecologically important behaviors like spatial navigation and action planning. Furthermore, it is unclear how far the results generalize to non-primate species.

We addressed these issues for Mongolian gerbils (*Meriones unguiculatus*) and designed a psychophysical task for time interval estimation of several seconds on a continuous range. The task was implemented in virtual reality (Thurley and Ayaz, 2017), which allows for the precise control of the behaviorally relevant variables.

First, we demonstrate the capability of gerbils to precisely measure and reproduce time intervals of several seconds. We also show that the gerbils’ responses display the regression effect, indicative of an error minimization strategy. Then, we present associated neural activity in gerbil medial prefrontal cortex (mPFC). This activity is composed of mixtures of responses including phasic activation and ramp-like firing patterns, response types well known from the interval timing literature (e.g. Gouvêa et al., 2015; Kim et al., 2013; Merchant et al., 2011; Mita et al., 2009; Paton and Buonomano, 2018). Since our task involves measurement and reproduction, i.e. the timing of an external event and of one’s own behavior, we could test how individual cells participated in both. To make the variety of responses accessible, we provide a comprehensive characterization of activity at the single neuron level and show that, despite the response heterogeneity, the mPFC population jointly measures and reproduces time intervals lasting several seconds. We find that distinct types of ramping neurons are necessary to explain the regression effect and thus for a mechanistic understanding of the neural basis of magnitude estimation behavior. This reveals that variables underlying cognitive functions may be encoded by mixed response types within a local neural population.

## Results

### Behavioral characteristics of time reproduction in gerbils

We trained gerbils to measure and reproduce the duration of time intervals lasting a few seconds. After the presentation of a black screen, the animals had to reproduce its duration by walking along a corridor in virtual reality. Intervals were randomly sampled between 3 and 7.5 s. Figure 1A-C details the apparatus and task. Gerbils exhibited typical magnitude estimation behavior (Petzschner, Glasauer, and Stephan, 2015). Across the range of stimuli, small stimuli were overestimated whereas large stimuli were underestimated, i.e. the regression effect. This effect was obvious in single sessions (Fig. 1D) and consistent across sessions and animals, as is displayed by the slope of the linear fits between stimuli and reproductions in a session being smaller than 1 (slope, Fig. 1E). The data also showed scalar variability, i.e. the standard deviation of reproductions increased with stimulus magnitude (inset in Fig. 1C). Variability (coefficient of variation (CV), Fig. 1E) matched values typically found in interval timing experiments and correlated with the strength of regression (Fig. 1F). Some sessions showed a general under- or overestimation in addition to the regression effect (BIAS, Fig. 1E).

**Figure 1.**
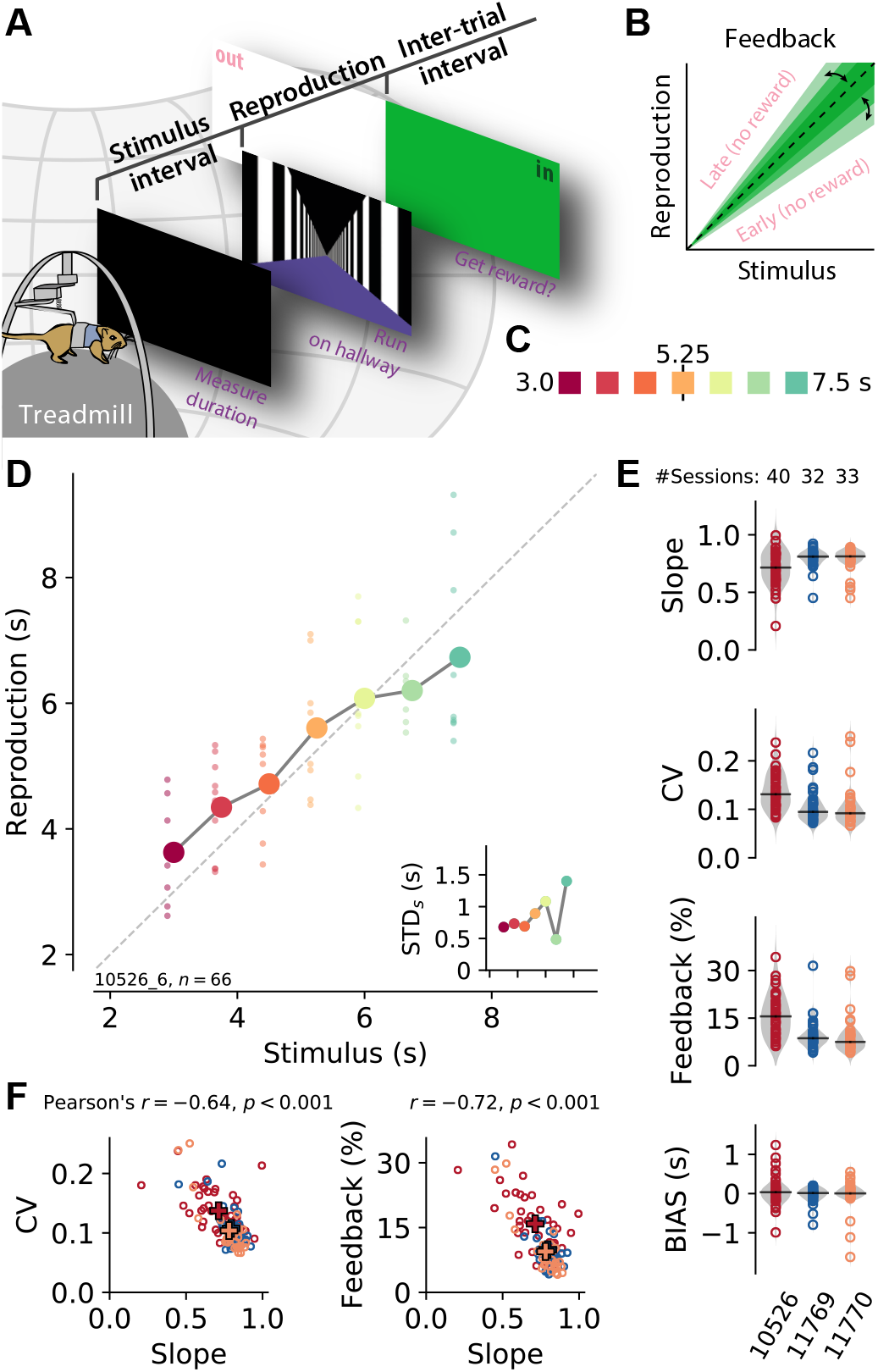
A time reproduction task for rodents. (A) Experimental apparatus and task. A gerbil was placed on top of a treadmill surrounded by a projection screen. Each trial started with a timed stimulus (black screen). The animal had to measure its duration and, when a virtual linear corridor appeared, reproduce the duration by walking. If the reproduction was close to the stimulus (“in”, i.e. a hit) a food reward was delivered and the entire screen was set to green for 3-4 seconds before another trial was initiated; for non-hits, screen color was set to white (“out”). (B) The feedback range was narrowed/widened after each in/out response. (C) Stimulus intervals were randomly sampled from a discrete uniform distribution with 7 values between 3 and 7.5 s. Colors identify stimulus duration and will be used throughout the paper. (D) Behavioral responses exhibited characteristic effects of magnitude estimation. Single reproductions (small dots) and their averages (large connected circles) showed the *regression effect* (data from one example session). *Inset:* Standard deviation increased with stimulus magnitude (*scalar variability*). Same *x*-axis as in the main panel. (E) Slope of the linear regression between stimuli and reproductions – quantifying the strength of the regression effect, with values closer to 1 meaning less regression –, coefficient of variation, average tolerance *k* of the feedback range, and average bias for each session sorted by animal (open circles). Colors identify animals. Gray violin plots illustrate the distribution of the population. A solid black line marks the median. (F) Slope negatively correlates with CV and feedback range across animals and sessions, indicating stronger regression effects with more variable responses. Open circles correspond to single sessions. Crosses mark averages for each animal. Color-code as in (E).

To keep the animals motivated, we gave them feedback on their reproduction performance (Fig. 1A). Although this feedback could be used to correct reproduction errors, we still observed the regression effect. Since we adapted the width of the feedback range after every stimulus (Fig. 1B), it could be used as an indicator of estimation precision, with a narrower feedback range corresponding to higher precision. The average width of the feedback range in a session indeed correlated negatively with the slope. In line with this, CV and slope also correlated negatively (Fig. 1F).

### Single cells differentially encode time during measurement and reproduction

We recorded a total of 1842 mPFC units over 105 experimental sessions from three gerbils. Visual inspection of the spiking responses, after sorting by stimulus and splitting into the two task phases “measurement” and “reproduction”, revealed a variety of response patterns that could underly time reproduction. Three examples are shown in Figure 2, further examples can be found in Figures S4-S6. The neuron in Figure 2A increased its activity during measurement, such that the firing rate by the end of the phase correlated with the stimulus interval (see Fig. 2D). During reproduction, this neuron displayed downward-ramping. However, here the change in firing rate correlated with the stimulus interval to be reproduced, such that for shorter intervals the firing rate decreased faster than for longer intervals (see also Fig. 2D). This effect was also observed in cells whose activity increased to a fixed level at the end of reproduction that was independent of the stimulus interval (Fig. 2C&D). Yet other cells constantly increased firing rate (Fig. 2B), similar to what we saw during measurement in the neuron in displayed in Figure 2A. We also found cells that responded at absolute times (Fig. S5) or relative to the reproduction interval, e.g. at its begin or end (Fig. S6A).

**Figure 2.**
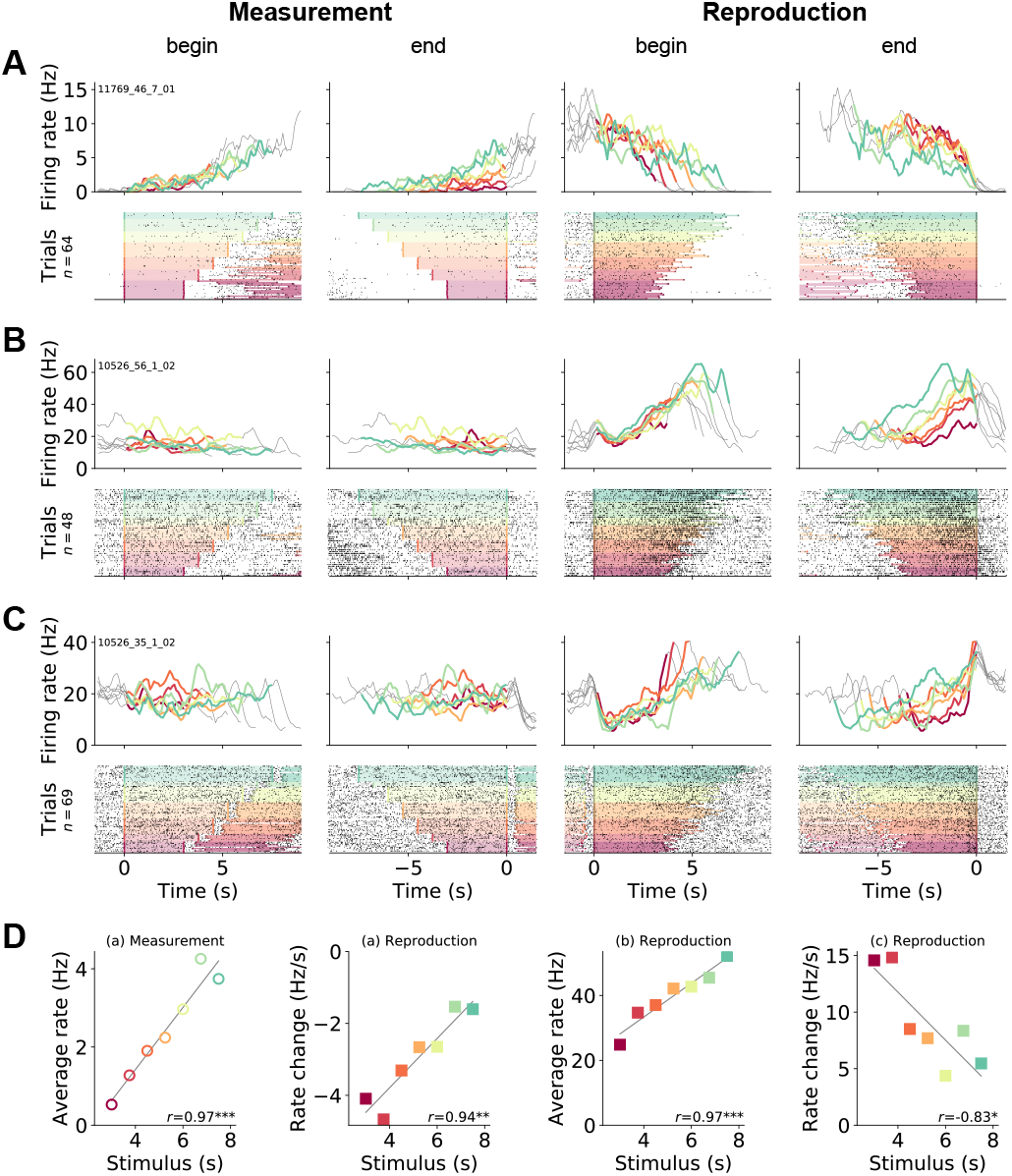
Gerbil mPFC neuron responses during time reproduction. (A) A cell that linearly increased its firing rate during measurement and ramped down to zero during reproduction. (B) A neuron that scaled its firing with the stimulus during reproduction and (C) a ramp-to-threshold cell. (A-C) Panels display spike raster plots sorted by stimulus (bottom) and corresponding spike density functions (SDF, top). Each column plots the data with different alignment, measurement begin and end, reproduction begin and end. Color identifies stimulus as in Fig. 1C. In the raster plots, black ticks are single spikes. For better visualization, we only plot half of the spikes (randomly chosen). Measurement or reproduction phases are delimited by underlayed color. The SDFs are colored in the respective task phase, outside they are displayed as thin gray lines. (D) Dependence of firing on stimulus duration in the example cells. Single markers show the average firing rate or change of firing rate at each stimulus. Open dots are used for data from measurement and filled for those from reproduction. Solid lines are linear fits. Pearson’s correlation coefficient and significance is given in the lower right corner. Above each panel, cell and task phase is indicated. The averages of firing rate and its change were calculated from the last half of the SDFs in the corresponding task phase.

Between measurement and reproduction, the neurons adapted their response patterns. We did not find a single cell that responded in the very same way in both task phases and essentially repeated its activity pattern from the measurement phase during reproduction – an observation that will be analyzed systematically below.

Since reproduction involved walking in our behavioral task, we also looked at dependencies between firing rate and running speed (Figs. S4-S6). In most example neurons, running speed modulation was weak and could not explain the observed firing patterns. Still, some neurons had an obvious speed modulation. Their response pattern was symmetrically rising and decreasing over the reproduction phase (Fig. S5G&H). We note that running speed as well as other unobserved behavioral factors may contribute to firing in some neurons and hence may explain differences in firing patterns between task phases. However, since we were interested in the collective action of the neuronal population, we did not investigate this further and also decided against excluding such neurons in the subsequent analyses.

### Single cell picture holds for the whole population

The different response patterns for single neurons were also obvious when visualizing the whole population. Some neurons ramped up at the end of measurement, others were active at the beginning and then decreased activity (Fig. 3A). During reproduction, a similar but more pronounced pattern emerged with up/down ramping and phasically active neurons (Fig. 3B).

**Figure 3.**
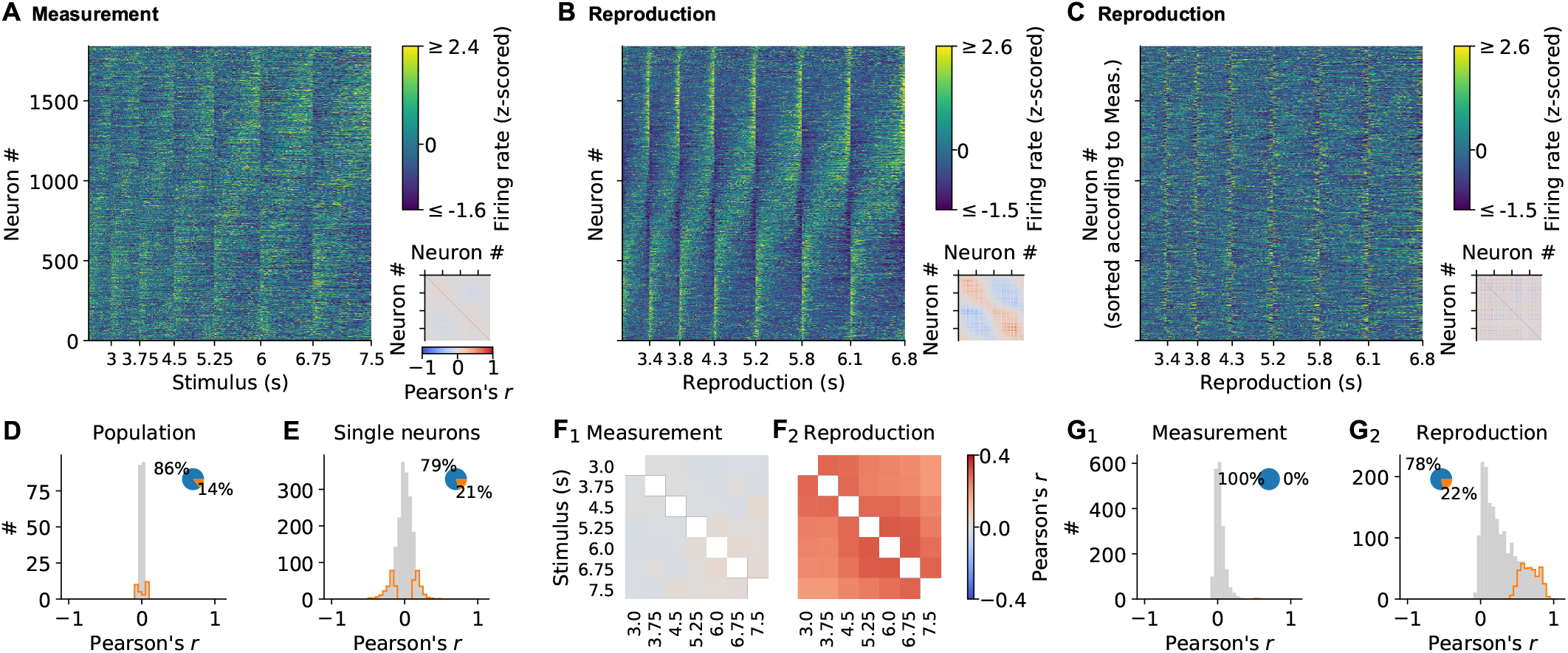
Single neuron and population dynamics in measurement and reproduction. (A) Normalized (z-scored) SDFs of all 1842 mPFC neurons for each stimulus interval during the measurement phase sorted by their timing within the intervals. *Small panel:* Matrix of pairwise Pearson correlations between all neurons. Diagonal entries not plotted. (B) Same as (a) but for the reproduction phase. (C) Same as (B) but with neurons sorted as in (A). (D) Population vector correlations in corresponding time bins of measurement and reproduction. (E) Pearson correlations between measurement and reproduction for single neurons. (F) Pairwise correlations of the whole population for different stimuli in measurement (F_1_) and reproduction (F_2_). Diagonal entries not plotted. (G) Distributions of average Pearson correlations of single cell activity for different stimuli in measurement (G_1_) and reproduction (G_2_). Histograms in (D,E&G) are displayed in gray with significant values delimited by an orange outline. Pie plots show significant (orange) and non-significant (blue) percentages.

Striking, however, were the activity differences between measurement and reproduction, indicating a state change in the population between both task phases. When individual cells were sorted in the same order for both measurement and reproduction, no global pattern was visible (cf. Fig. 3B&C). Also, correlating population vectors in corresponding time bins for measurement and reproduction yielded only low values (Fig. 3D). For single cells, however, the correlation between task phases was larger and significant in about 20% of the cells (Fig. 3E). Note that this does not indicate a precise correspondence between the activity profiles for measurement and reproduction but rather that a neuron was active in both phases. See, e.g., the neuron in Fig. 2A, which shows a negative correlation between task phases.

Neural activity was similar for different stimulus magnitudes during reproduction. On the one hand, population activity correlated for different stimuli (Fig. 3F_2_); on the other hand, single neuron activity correlated across different stimuli in about 20% of the neurons (Fig. 3G_2_). During measurement, correlations across stimuli were absent (Fig. 3F_1_&G_1_).

The above picture remained, when we split activity into odd and even trials. Activity was largely similar arguing for stable neural activity throughout a session. However, during measurement, data was more noisy in agreement with the weak correlations we observed in this task phase (Fig. S7).

### Scaling of neuronal activity with duration

The activity of the prefrontal neurons we recorded appeared to scale with stimulus duration (Fig. 3A&B), which was reminiscent of findings in interval timing studies for striatum (Mello, Soares, and Paton, 2015) and prefrontal cortex (Wang et al., 2018; Xu et al., 2014).

To examine stimulus-related scaling in our data set, we followed the approach of Mello, Soares, and Paton (2015). We first calculated for each cell the center of mass (COM) of the SDF at every stimulus duration. The COMs were, in particular during reproduction, widely distributed and tiled the time intervals. For comparison, we simulated SDFs with no temporal modulation (“noise”). The corresponding COMs clustered around the center of the intervals and significantly differed from those for the recorded SDFs (*p* < 0.001 in all cases, Kolmogorov-Smirnov test; Fig. 4A). Interestingly, during reproduction, for neurons with a COM at the begin and end of an interval, activity correlated more strongly across different stimuli (Fig. 4B, cf. Fig. 3G_2_). This indicates that, in particular for neurons with increased firing at the border of the interval, activity depended on the temporal stimulus.

**Figure 4.**
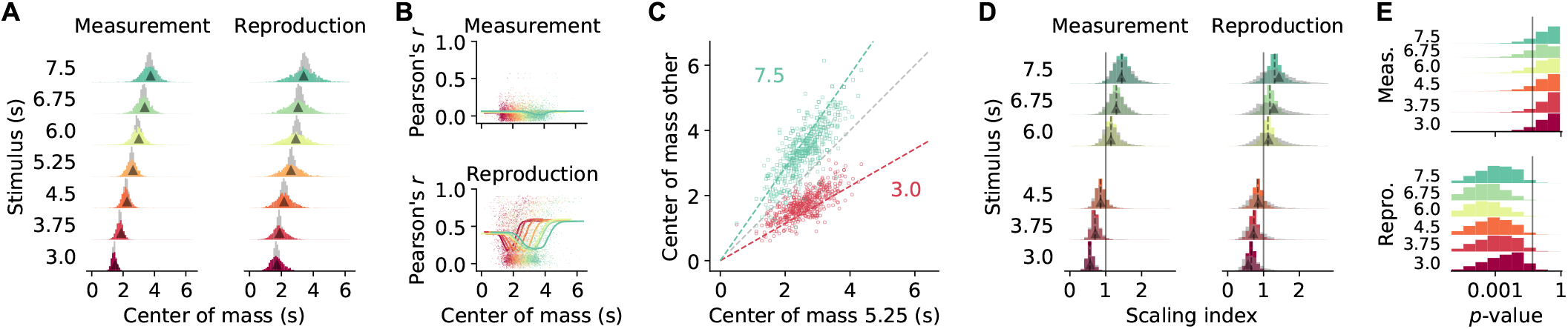
Temporal scaling. (A) Distributions of center of mass for all 1842 mPFC neurons (colored histograms). Grey histograms give distributions of center of mass for noise SDFs. Arrow heads mark the middle of the stimulus interval and of the average reproduced interval, respectively. (B) During reproduction, the average Pearson correlations of single cell activity for different stimuli (cf. Fig. 3F) were larger for neurons with center of mass at the begin and end of an interval. Dots give single cell data, solid lines are moving averages. Arrow heads in the lower panel mark the middle of the average reproduced interval. (C) Center of mass of each cell for 5.25 s against 3 s (red) and 7.5 s (green) during reproduction. Colored dashed lines mark the predicted ratio; gray dashed line corresponds to no change. (D) Distributions of scaling indices for all cells, i.e. center of mass of the SDF at every stimulus divided by the center of mass for 5.25 s. Solid vertical line marks one. Dashed lines mark the medians and arrow heads the ratio between 5.25 s and the respective other stimulus. Grey histograms give distributions of center of mass for shuffled control SDFs. (E) Bootstrapped distributions of *p*-values for Kolmogorov-Smirnov tests between scaling indices for recorded and shuffled SDFs. Solid vertical line marks 5%. Logarithmic *x*-axis.

As a second step, we calculated scaling indices by dividing every cell’s COMs by the corresponding COM at the 5.25 s stimulus, i.e. the mean of the stimulus distribution. These scaling indices theoretically range between 3.0/5.25 ≈ 0.57 and 7.5/5.25 ≈ 1.43. Indeed scaling indices were close to the theoretical values during both measurement and reproduction (Fig. 4D; see also Fig. 4C). During reproduction, scaling indices were even larger or smaller than the prediction in a manner consistent with the regression effect, i.e. smaller durations had larger scaling factors and vice versa.

Finally, we obtained scaling indices after shuffling cell identities across stimuli to test whether the scaling indices from the recorded data were in fact the result of a meaningful modulation. Only for the reproduction phase, the indices from shuffling were more widely distributed than the data; for the measurement phase a difference was not clearly visible (Fig. 4D). Statistical testing, however, identified significant effects in both phases (*p* < 0.01 in all cases, Kolmogorov-Smirnov test). This statistical result was likely due to the large number of samples contained in the data set. Therefore, we performed bootstrapped Kolmogorov-Smirnov tests on 10% of the data over 10.000 runs. The *p*-values obtained lay below 5% for almost all cases during reproduction. During measurement only about 30% of the bootstrapped Kolmogorov-Smirnov tests were significant (Fig. 4E).

Taken together, this indicates activity-dependent temporal scaling of prefrontal single cell responses in the reproduction phase alluding to a potential role in time encoding and the regression effect. Scaling during measurement was not different from what would be expected of neural activity that unfolds over time. Note, that the results are in line with the non-/significant correlations of activity across stimuli for measurement and reproduction, respectively (cf. Fig. 3F&G).

### Population activity shares similar components for measurement and reproduction

The activity differences between measurement and reproduction revealed above argue against a magnitude estimation mechanism just at the single cell level. Therefore, we examined the collective properties of prefrontal neurons by decomposing the population activity into its principal components (PCs). We used demixed PCA a form of principal component analysis (PCA) that allowed us to separate time course-related from stimulus-dependent contributions to neural activity (Kobak et al., 2016).

Despite dissimilar responses of single cells in measurement and reproduction, PCs looked very much alike for both task phases (Fig. 5A&B). The time course-related PCs were ramp-like (PC 1) or had a comparable non-monotonous shape (PC 2&3). The strongest stimulus-dependent PC was constant over time and had amplitudes that were ordered by stimulus (see bottom right panels in Fig. 5A&B). Moreover, the contribution of the PCs to the single cell responses (PCA scores) correlated for PC 1 and stimulus PC 1 in both measurement and reproduction, indicating a link between ramp-like and stimulus-dependent activity (Fig. 5C, see also Fig. S10). During reproduction but not during measurement, stimulus PC 1 also correlated with PCs 2&3, arguing for a richer representation of the to-be-reproduced compared to the to-be-measured time interval. Note that, in the measurement phase, PCs 2&3 in general contributed very little to explaining population activity (Fig. S8).

**Figure 5.**
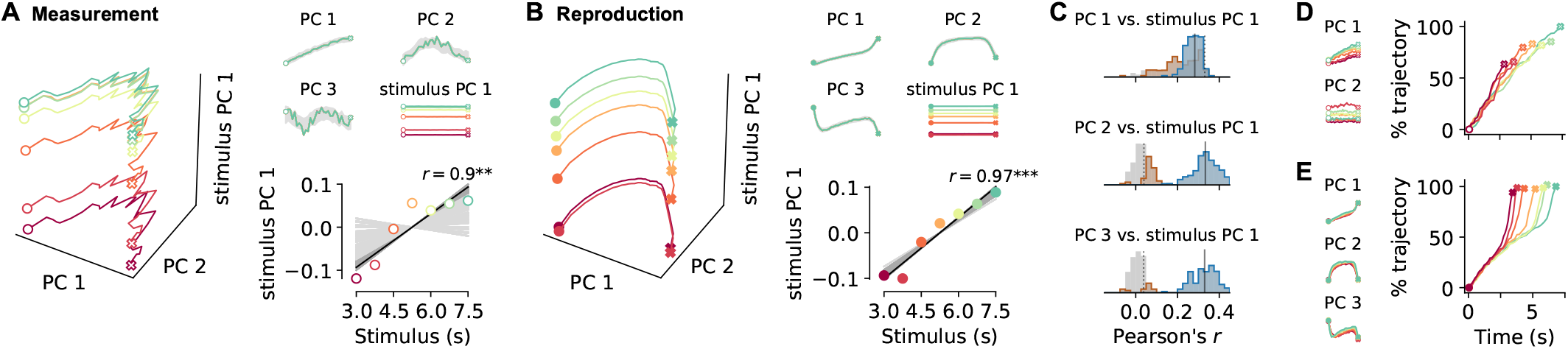
Decomposition of population activity. Demixed PCA yields separate principal components for the time course during a trial (PC 1-3) and for each stimulus (stimulus PC 1). (A) PCs for measurement and (B) for reproduction. The small panels display individual PCs at each stimulus scaled to the length of the interval. Note that PC 1-3 for different stimuli lie on top of each other, demonstrating perfect demixing. Stimuli are colored as in previous figures. Circles and crosses mark interval start and end. Open symbols are used for measurement and filled for reproduction. The PCs were calculated for the whole data set and gray-shaded areas delimit the standard deviation of results from bootstrapping with 10% of the cells. Bottom right panels in (a) and (b): Correlation of stimulus PC 1 with the stimulus. Black solid line is a linear fit and gray lines are fits for the bootstrap samples; dark gray significant, light gray non-significant cases. (C) Distributions of Pearson’s correlation coefficients between PC 1-3 and stimulus PC 1 from bootstrap samples. Significant correlations are colored in brown (measurement) and blue (reproduction). Dotted (measurement) and solid (reproduction) black lines mark correlations coefficients for the whole data set. (D, small panels) PCs for conventional PCA during measurement scaled to the length of the interval for each stimulus. Note that, in contrast to demixed PCA, conventional PCs contain both time course-related and stimulus information. (D, large panel) Time course of PC 1 for conventional PCA at each stimulus during measurement. The percentage of covered trajectory is given with respect to PC 1 for the 7.5 s stimulus. (E) Same as (D) for reproduction.

Recent work showed that when different time intervals are estimated collective neural activity evolves along very similar trajectories, but which final state is reached depends on the duration of the interval. When a time interval is reproduced trajectories of similar length are found for different stimuli but now the speed of progression decreases with duration (Gouvêa et al., 2015; Remington et al., 2018; Sohn et al., 2019; Wang et al., 2018). Since we separated time course-related from stimulus information through the demixed PCA, we could not see such stimulus-dependent effects directly. We therefore also applied conventional PCA to our data. The strongest PCs from conventional PCA were similar in shape to those from demixed PCA but were – as expected – influenced by the stimulus (Figs. 5D&E and S9). Indeed, in the measurement phase, neural activity followed trajectories whose length depended on stimulus duration. During reproduction, trajectories had similar length. However, activity along those trajectories accelerated over an interval resulting at higher average speed for stimuli of shorter duration. Note that this effect corresponds to the temporal scaling described in the previous section.

It is possible that the population responses during measurement and reproduction are not actually collective phenomena but simply the result of pooling single neurons. To test this, we compared population activity to surrogate data that preserve stimulus tuning of single neurons, correlations of single cell firing rates across time, and pairwise correlations between neurons (Elsayed and Cunningham, 2017). During measurement, PC 1 of the original data was larger than expected, suggesting that collective activity added to representing ongoing time; similarly, PC 3 was larger than expected during reproduction (Fig. S11). Stimulus PC 1 was larger than expectation for reproduction and at the lower end of what was expected during measurement, suggesting that the population as a whole did contribute to stimulus representation. Collective activity thus contributed to the representation of ongoing time and the representation of the time stimulus.

### Heterogeneous prefrontal activity can be categorized into a few response types

To extract response types from a set of neurons, usually, specific tests must be designed, which are based on predefined knowledge of the responses of interest. To circumvent such rigid preselection, we used the demixed PCA results for categorizing neurons into different response types.

Single cell activity patterns can be decomposed into linear mixtures of different PCs. We focused on the contributions of the strongest PCs, which were time course-related PC 1 and stimulus PC 1 during measurement and PCs 1-3 and stimulus PC 1 during reproduction (Fig. S8). As we already mentioned above, the PCA scores for single cell responses were correlated for these PCs (Fig. 5C), pointing to a potential utility for capturing single cell responses. For instance, mixing the ramp-like PC 1 and stimulus PC 1 with its constant responses ordered by stimulus, one can describe the activity during the measurement phase of cells like in Fig. 2A. Similarly, mixtures of the different time-modulated profiles from PCs 1-3 and stimulus PC 1 can capture various response patterns during reproduction (Fig. 6A).

**Figure 6.**
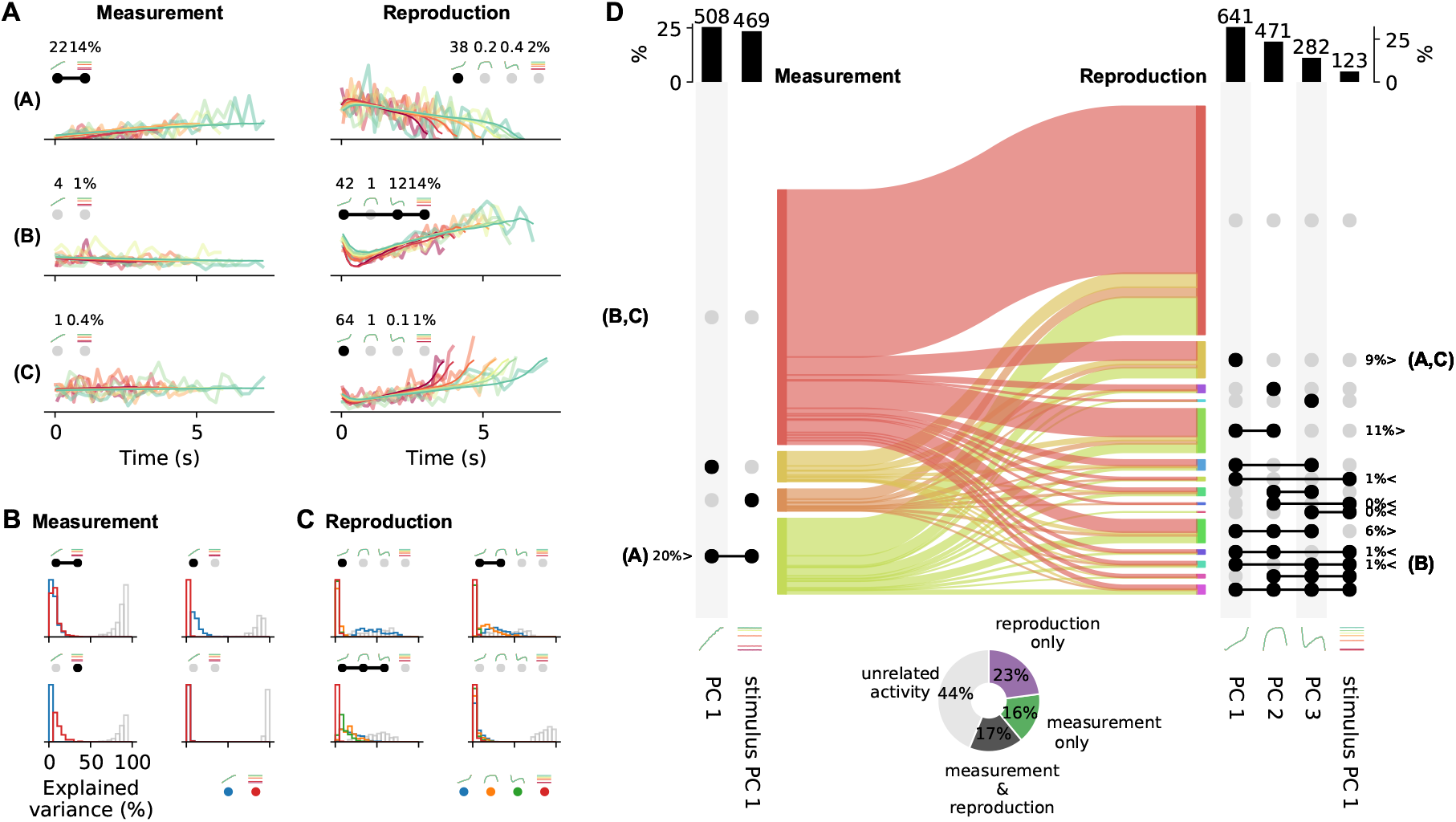
Single cell activity and their contribution to the population response change from measurement to reproduction. (A) Reconstructions (thin lines) of the firing patterns (SDFs, thick faint lines) of the example neurons from Fig. 2 (identified by bold letters; note that here SDFs were not smoothed). Markers show percent explained variance for each PC and illustrate the response type according to the categorization (see also (D)). For instance, the neuron in the first row is reconstructed by the linear combination of PC 1 and stimulus PC 1 during measurement and with (negative) PC 1 during reproduction. (B) Distributions of explained variance contributed to single cell responses by PCs during measurement. Variance explained by PC 1 is displayed in blue and by stimulus PC 1 in red. The remainder (i.e. variance unexplained) is displayed as gray open bars. Each panel displays distributions for cells belonging to one of the four possible response categories visualized by the markers above the panel. (C) Same as (B) for the reproduction phase. Here the coloring is PC 1-3, blue, orange, green; stimulus PC 1, red. Distributions are displayed for the four categories with the most cells. (D) The main plot displays the numbers of cells with activity patterns that can be reconstructed by PCs. For measurement (left) PC 1 and stimulus PC 1 were considered and for reproduction (right) PC 1-3 and stimulus PC 1. Dark dots indicate the contribution of a PC to a response category. Percentages are only given for categories that contain significant numbers of cells compared to shuffled data. The signs < and > indicate if the number is smaller or larger than for shuffled data. Zero percentages mean less than 1%. Bold letters correspond to the examples from (A). Sankey diagram in the center illustrates the transition of cells between categories from measurement to reproduction. Bar graphs at the top show percentages and cell numbers with contributions of each PC. Pie chart at bottom shows percentages of cells active during the task phases determined from the reconstructions displayed in the main plot.

Combinations of PC 1 and stimulus PC 1 yield three possible response categories in the measurement phase: activity explained by (1) PC 1 only, (2) stimulus PC 1 only, or (3) both PCs. For each neuron a category was selected based on the angle between its PC 1 and stimulus PC 1 scores (Fig.S10C). We only included cells with large demixed PCA scores (explained variance) in this analysis (see also Fig. S10). Cells with small scores were categorized as “unrelated activity”, giving a fourth response category. Likewise, 15 different categories were defined for the reproduction phase from combinations of PCs 1-3 and stimulus PC 1 plus a sixteenth category for “unrelated activity”.

Categorisation provided meaningful representations of single cell responses as reflected in the variance explained by the contributing PCs (Fig. 6B&C). For instance, large parts of variance were explained by PC 1 for neurons in the category “PC 1 only” during measurement; contributions from stimulus PC 1 were marginal. In the stimulus PC 1 only-category, the situation was reversed, and for the mixed PC 1 & stimulus PC 1 category contributions by both PCs matched (Fig. 6B). However, only less than 50% of variance could be explained by the two PCs in the measurement phase. The PCs used for reconstructing during the reproduction phase better captured the activity; leaving less variance unexplained (Fig. 6C). In particular, for the PC 1 only-category often cell activity could be explained to more than 50% by PC 1.

Cells that could be described by PC 1 had a ramp-like response profile. During measurement, about a quarter of the cells showed such ramping activity. These cells included ramping to stimulus-dependent levels like the neuron in Fig. 2A and ramping that did not depend on stimulus. The other cells represented the stimulus in a different way or showed activity unrelated to the task during measurement (67%); Fig. 6D.

During reproduction, about 10% of the neurons showed ramping activity (PC 1 only), e.g. ramp-to-threshold cells where the rate of ramping decreased with stimulus magnitude (cf. examples in Figs. 2B&C, S4B&C). Similar fractions were explained by mixing PCs 1 and 2 and by a mixture of PCs 1-3. Only 1% was explained by ramps that encoded stimulus (i.e. PC 1 & stimulus PC 1). However, counting all combinations of stimulus PC 1 with PCs 1-3 showed that about 5% of the neurons combined ramps with stimulus-dependent firing. In total, 35% of the cells contained a ramping component. These cells overlapped largely with those whose activity was significantly correlated across stimuli (not shown, cf. Fig. 3G_2_). In Figures S4-S6 further examples for the different categories can be found.

The categorization analysis strengthened the picture of changing response patterns between measurement and reproduction we already observed at several instances above. Cells switched response types between task phases but with no specific pattern as becomes obvious from the flow diagram in Fig. 6D.

Separating cells with small scores from those with large ones, we could estimate the number of cells active during the task phases. Close to a fifth of the cells fired action potentials in both phases, 16% were only active during measurement and 23% only during reproduction; 36% of the cells did not contribute substantially in either phase (pie chart in Fig. 6D).

### Time encoding is governed by different response types

To understand how the prefrontal neuronal population represents time, we used simple linear regression to decode time from our recorded responses (see Methods). We began by decoding time from the responses of all neurons during the reproduction phase. Time readouts were accurate in the first seconds, however, by the end of the phase a regression effect was visible with over- and underestimation at short and long stimuli, respectively (Fig. 7A). The effect was due to neurons with large PCA scores. Cells with small scores led to decoding estimates more similar to shuffled data, where decoded time was almost constant throughout the interval, such that real time was overestimated initially and underestimated at the end of reproduction. This picture was even pronounced when we decoded time from pure noise (Fig. S14).

**Figure 7.**
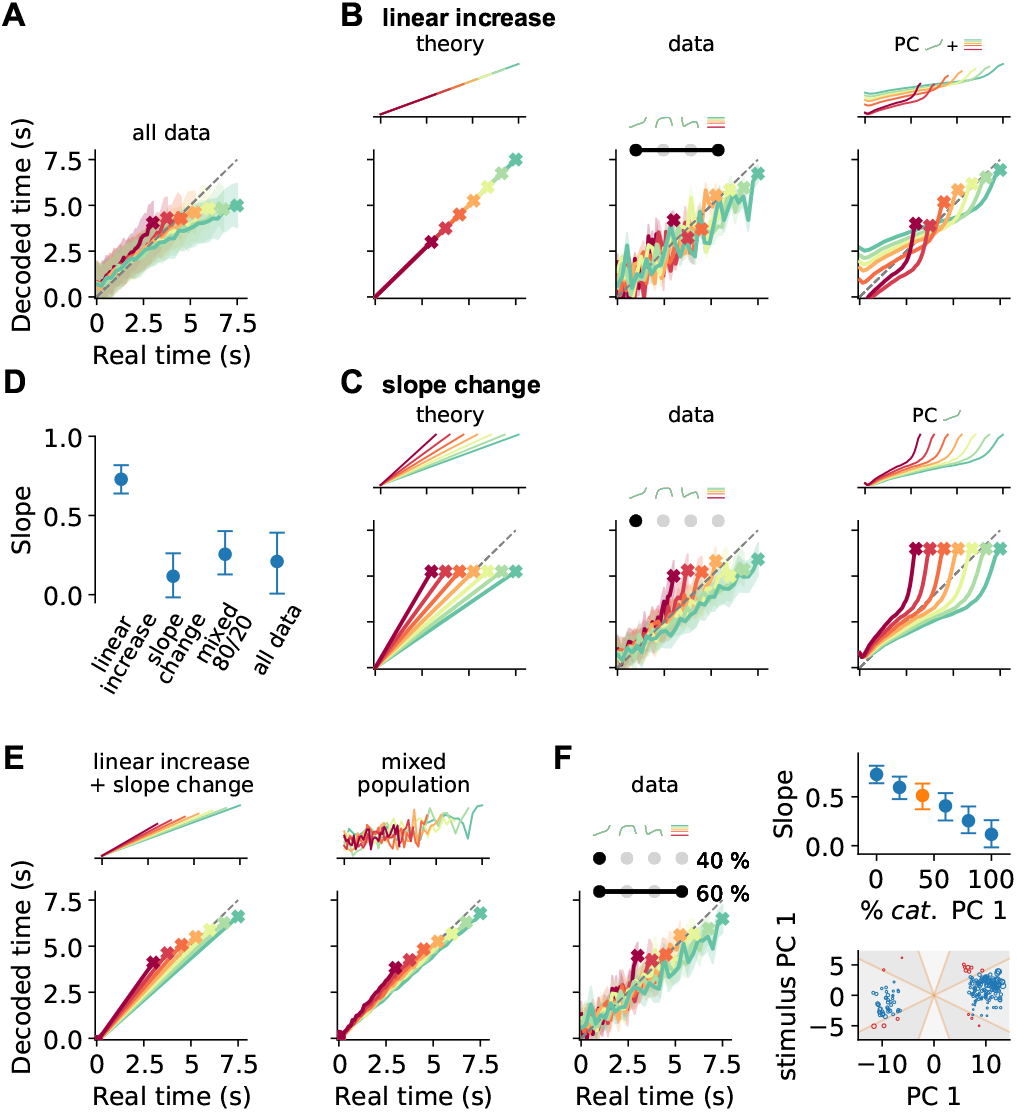
Decoding elapsed time from prefrontal population activity. (A) Elapsed time (avg±std from bootstrapping) decoded from the responses of all neurons recorded during the reproduction phase. As before, color identifies the stimulus. Crosses mark final values. A strong regression effect is visible. (B) Time decoding with neurons that ramp to stimulus-depended levels but with the same slope and (C) with ramp-to-threshold cells. Left panels illustrate the theoretical prediction for decoded time and example neuronal activity above. Middle panels plot decoding results using only neurons from the respective response category. Right panels give results for decoding using the corresponding components from demixed PCA. As displayed in the top panels, these components are PC 1 and stimulus PC 1 in (B); and PC 1 only in (C). (D) Median slopes of the linear regression between the final values of real and predicted time for decoding from data in (A) and for a mixture of 80% slope changing and 20% linear increasing cells; cf. (C). Error bars delimit interquartile ranges (from bootstrapping). (E) Mixtures of linear increasing activity and slope changes explain behavioral regression effects. Theoretical predictions for mixing both responses in single neurons (left) or as two different response types across a population (right). For the second case, a neuron with noisy linear increasing activity is display as an example. (F) Decoding results for a mixed population of 40% slope changing and 60% linear increasing cells. The cells were sampled at these fractions in each bootstrapping run from the response categories we identified in our recorded data. The upper right panel shows the regression slope (cf. D) for different fractions of slope changing cells. The orange marker corresponds to the example in the left panel. The lower right panel displays the PCA scores of the cells from the linear increasing (red; PC 1 + stimulus PC 1) and slope changing categories (blue; PC 1 only). The size of the marker illustrates the decoder weight *β* for that cell. See also Fig. S10C.

We wondered what response types could mediate the regression effect and simulated a few stereotypical cases. Decoding time from neurons that ramp with same slope to stimulus-dependent levels (linear increasing neurons) show precise time representation but no regression effect (Fig. 7B). Whereas, ramp-to-threshold cells that change slope but reach same activity levels by the end of an interval can only encode the mean of the stimulus distribution and result in maximal regression (Fig. 7C).

We got similar results, decoding time from the response categories matching the theoretical cases (cf. Fig. 6). For the response type combining time-dependent PC 1 with stimulus PC 1, time reproduction displayed only a weak regression effect (Fig. 7B) and ramp-to-threshold responses (PC 1 only) displayed very strong regression (Fig. 7C); see also Fig. 7D. Decoding time directly from the demixed PCs gave similar results (Fig. 7CD).

Regression effects in our behavioral data lay in between the extremes yielded by the two response categories (cf. Fig. 1E). This discrepancy made us think about another solution to the question of how regression effects may arise. In theory, combining ramping to stimulus-dependent levels with slope-changes by stimulus also generates regression effects (Fig. 7E). Such a combination can be implemented either with mixed response patterns within one neuron or as a mixture of response types across a neuronal population. To find out which of the two scenarios underlies our data, we looked at the distribution of PCA scores for the cells in the linear increasing and the slope-change categories. These scores were broadly distributed and also the corresponding decoding weights did not reveal a particular structure, indicating time coding comprising different response types (bottom-right panel in Fig. 7F). To further test this possibility, we mixed responses of recorded cells from both categories at different fractions. Decoding not only yielded regression effects, but the strength of regression could be manipulated by the relative shares of both response types. The more slope-change cells were present in the population, the stronger was the regression effect (Fig. 7F).

Another response type that has been often connected to timing are phasically active cells. We therefore wondered how ongoing time would be encoded by such neurons. Constructing again theoretical stereotypes, we saw that neurons that are active relative to the stimulus interval can only encode a single time point. In contrast, a population of neurons that are phasically active at different absolute times tiling the whole interval would provide an accurate time representation (Fig.S15). Since neither relative nor absolute timing neurons by themselves predict regression effects as found in the behavioral responses, again a mixture of both response types would be required. Matching these theoretical results to our own recordings turned out to be less straightforward. We recorded phasically active cells (cf. Fig. S5) but this subset of cells did not completely tile the whole time intervals. However, this prerequisite is necessary to seriously attempt to reproduce the theoretical predictions.

Finally, we examined whether time could also be decoded from the neuronal responses in the measurement phase. Here, decoding was imprecise with overestimation at the begin and underestimation at the end of the interval (Fig. S16) – a picture that also appeared for shuffled and noisy data (cf. Fig. S14) and matches the less pronounced time signalling across the population during measurement, which we have found above in several places. A general underestimation was also found when we used only neurons in the PC 1 + stimulus PC 1 category, indicating their foremost influence on the collective readout. Interestingly, when we decoded time from the cells in the PC 1 only-category, a regression effect was also seen during measurement (rightmost panel in Fig. S16), suggesting an impact of previous stimuli (prior knowledge) on the activity of these neurons already during stimulus measurement and not just during reproduction.

## Discussion

We investigated the neural basis of magnitude estimation and analyzed neural correlates from rodent mPFC in a novel interval timing task. We showed that Mongolian gerbils (*Meriones unguiculatus*) are able to measure and reproduce durations lasting several seconds. To allow the gerbils to respond in a natural way, we used walking as a response (Meijer and Robbers, 2014). The task was implemented in a rodent virtual reality system (Thurley and Ayaz, 2017), which (1) gave us the use of a treadmill, (2) prevented landmark-based strategies for task-solving and (3) decoupled time from distance, such that the task could not be solved by path integration.

The rodents’ behavior exhibited typical characteristics of magnitude estimation, including the regression effect, i.e. the overestimation of small and underestimation of large stimuli (also known as regression to the mean, central tendency, or Vierordt’s law; Hollingworth, 1910; Petzschner, Glasauer, and Stephan, 2015; Shi, Church, and Meck, 2013; Vierordt, 1868). Neural activity in gerbil mPFC correlated with and likely contributed to magnitude estimation behavior. Prefrontal neurons displayed various firing characteristics, which could be grouped into a number of representative categories. Single cell firing patterns differed between measurement and reproduction. For those cells that participated in both task phases, their activity profiles, although sometimes correlated, never matched between task phases; see e.g. the ramp-type neurons in Figures S4A,D&S6D. Moreover, activity changes between task phases were not coordinated across neurons, leading to low population vector correlations and no specific transition patterns between response types. Linear decomposition of the population activity, however, revealed state-space trajectories that were only slightly modulated during measurement and reproduction. This indicates that – despite the response heterogeneity within and between cells – the prefrontal population similarly encoded time in both task phases. Such effects on low-dimensional population activity in connection to changes of behavioral and cognitive state have been described for attention and task engagement (Engel and Steinmetz, 2019).

Neural correlates of interval timing in the range of seconds have been found in several brain areas (Issa et al., 2020; Merchant, Harrington, and Meck, 2013; Paton and Buonomano, 2018) including prefrontal cortex (Emmons et al., 2017; Genovesio, Tsujimoto, and Wise, 2006; Kim et al., 2013; Tiganj et al., 2017; Xu et al., 2014), pre-/supplementary motor cortex (Merchant et al., 2011; Mita et al., 2009), hippocampus (MacDonald et al., 2011), entorhinal cortex (Heys and Dombeck, 2018), and striatum (Bakhurin et al., 2017; Emmons et al., 2017; Gouvêa et al., 2015; Mello, Soares, and Paton, 2015). What distinguishes our experiments from previous studies is twofold: (1) We tested time intervals on a continuous range. (2) We combined timing of an external event (measurement phase) and timing own behavior (reproduction phase), linking sensory and motor timing (Paton and Buonomano, 2018). Our study is therefore conceptually different from the fixed interval or discrimination tasks that have been used in most of the above timing studies. Some studies with monkeys used tasks comparable to ours but focused on intervals lasting only hundreds of milliseconds (Jazayeri and Shadlen, 2015; Sohn et al., 2019). The neural activity in primate parietal and frontal cortices observed in these studies is surprisingly similar to what we found in rodent mPFC. This is especially interesting since we tested timing of several seconds and neural dynamics typically act on much shorter time scales.

The mPFC responses we recorded during the reproduction phase are reminiscent of the neural correlates of self-initiated behavior found in rat secondary motor cortex by Murakami et al. (2017, 2014). The reproduction phase in our task also involves self-initiated behavior (i.e. to stop walking). Murakami *et al*. found ramp-to-threshold cells similar to the one in Figure 2C. However, in contrast to our findings, they reported the absence of such responses in mPFC (Murakami *et al.*, 2017). This discrepancy may be due to the different tasks involved: waiting for a signal appearing after a random interval in their experiments vs. responding after a previously measured interval in our case. Ramp-to-threshold responses were also reported in monkey pre-/supplementary motor cortices (Merchant et al., 2011; Mita et al., 2009) and as a population pattern in lateral intraparietal cortex (Jazayeri and Shadlen, 2015) during time (re-)production, demonstrating their ubiquitous presence in self-initiated behaviors. Note, that we also recorded negative ramp-to-threshold, i.e. ramp-down, responses (Fig. 2A). In addition, we observed linear increasing neurons (neurons that ramp to stimulus-dependent levels at the same slope, e.g. Fig. 2B) that may serve as integrators of time information provided by sequentially-activated, phasically responding cells (Fig. S5). Again, both types of neurons have been reported in other timing tasks (Genovesio et al., 2016; Gouvêa et al., 2015; Kim et al., 2013; Merchant et al., 2011). Responses with a linear ramping component (comprising both linear increasing and ramp-to-threshold) have been shown to be important for precise time decoding and to underly low-dimensional population dynamics (Cueva et al., 2020).

Our time decoding analysis revealed that ramp-to-threshold and linear increasing neurons by themselves are not capable of explaining the regression effect; rather the combination of both response types is necessary in either single neuron activity or mixed across a population of neurons. This view is compatible with theoretical models of interval timing (Simen et al., 2011) and magnitude estimation (Thurley, 2016). Similarly, distributing relative and absolute timing cells across a neuronal population may yield the regression effect. However, since ramp-like responses were prominent throughout our data set, we consider the first option more likely. Mixed response types are in general important in cognitive tasks (Rigotti et al., 2013) but have also been reported during spontaneous behavior (Stringer et al., 2019). Coding of variables related to cognitive functions like choice and task engagement is often distributed across brain regions (Steinmetz et al., 2019). Although, we only recorded in one brain region and cannot comment on the distribution across the brain, our findings suggest that a local distribution of response types may also underly cognitive functions.

Temporal scaling appears to be a general feature of timed computations in the brain as it has been described in various brain regions (Mello, Soares, and Paton, 2015; Wang et al., 2018; Xu et al., 2014). It has also been demonstrated to be important for time-coding in neural network models (Bi and Zhou, 2020). In our experiments neuronal activity scaled and changed speed in relation to the stimulus duration in the reproduction phase only. Here, animals had to actively generate timed behavior (motor timing); in contrast to the measurement phase (sensory timing), where due to the randomization stimulus duration could not be determined in advance.

Temporal scaling and speed-dependence of neural dynamics corresponded to the regression effect we observed in the behavioral data. Bayesian models (Jazayeri and Shadlen, 2010; Petzschner, Glasauer, and Stephan, 2015) as well as other approaches (Bausenhart, Dyjas, and Ulrich, 2014; Thurley, 2016) have demonstrated that the regression effect may be a strategy to minimize behavioral errors. Bayesian models fuse probability distributions of the current stimulus estimate and prior knowledge. The neural representations of these probability distributions and the mechanism underlying the probabilistic computations has yet to be determined. An interesting solution was recently proposed based on recordings from monkey frontal cortex by Sohn et al. (2019). While the animals measured time intervals in a task very similar to ours, frontal cortex activity followed low-dimensional curved state-space trajectories. These curved trajectories can be interpreted as a compressed non-linear representation of time, which when read-out appropriately during reproduction, can explain regression effects seen in behavior. Our demixed PCs also showed curved trajectories during measurement (Fig. 5A); however, trajectories from conventional PCA did not comprise such curvatures (Figs. 5D&E and S9A).

Our results point to another solution, namely the regression effect as the consequence. of mixed neural responses comprising two different types of ramping. When these types are implemented in different neurons and distributed across the population, the amount of noise in the system determines the strength of the regression effect. Without noise, magnitude encoding would be dominated by neurons with linear increasing activity at constant slope and no regression effect could emerge. If activity is noisy, neurons with stimulus-dependent slope contribute and the regression effect appears. In fact, the amount of noise (uncertainty about the current stimulus) determines the impact of either response type and thus the balance between current stimulus estimate and prior knowledge.

State-space trajectories for different stimuli were well separated during measurement (Fig. 5A). Since stimulus duration is unknown at the beginning of the measurement phase, stimulus-dependent trajectories are likely due to prior expectations about the stimulus. Small stimuli are typically followed by larger ones and vice versa, which may bias neural responses and behavioral estimates accordingly. Such sequential effects are known in magnitude estimation (Bausenhart, Dyjas, and Ulrich, 2014; Petzschner, Glasauer, and Stephan, 2015; Thurley, 2016). Interestingly, when we only included ramping cells for time decoding a regression effect was also seen during measurement (Fig. 7D), implying that previous stimuli affect the current measurement. During reproduction, trajectories were also ordered by stimulus (Figs. 5B&S9B), which is considered an indication that cortical dynamics are adjusted for (re-)producing different time intervals (Remington et al., 2018).

The present work provides first insight into the neural substrate of magnitude estimation, including the regression effect and error minimization, in rodents. A thorough characterization of mPFC responses during time-interval reproduction allowed us to show that only mixed responses in either single cells or distributed across a local population of neurons can explain the regression effect. Adjusting the relative fractions of response types one can parameterize the strength of the regression effect and thus the fusion of stimulus estimate and prior knowledge. To resolve the specifics of the underlying neural computations will be an important direction for future research.

## Methods

### Animals

The experiments in this study were conducted with three female adult Mongolian gerbils (*Meriones unguiculatus*) from a wild-type colony at the local animal house (referred to by ids 10526, 11769, and 11770). Training started at an age of at least four months. The gerbils were housed individually on a 12-h light/dark cycle, and all behavioral training and recording sessions were performed in the light phase of the cycle. The animals received a diet maintaining them at about 85-95% of their free feeding weight. All experiments were approved according to national and European guidelines on animal welfare (Reg. von Oberbayern, District Government of Upper Bavaria; reference numbers: AZ 55.2-1-54-2532-10-11 and AZ 55.2-1-54-2532-70-2016).

### Behavioral experiments

#### Experimental apparatus

Experiments were done on a virtual reality (VR) setup for rodents (Fig. 1A). For a detailed description see Thurley et al. (2014). In brief, the setup consists of an air-suspended styrofoam sphere that acts as a treadmill. On top of the sphere the rodent is fixated with a harness that leaves head and legs freely movable. Rotations of the sphere are induced when the animal moves its legs. The rotations are detected by infrared sensors and fed into a computer to generate and update a visual virtual scene. The scene is displayed via a projector onto a projection screen that surrounds the treadmill. We used Vizard Virtual Reality Toolkit (v5, WorldViz, http://www.worldviz.com) for real-time rendering; the virtual environment was designed with Blender (v2.49b, http://www.blender.org/). Animals were rewarded with food pellets (20 mg Purified Rodent Tablet, banana & chocolate flavor, TestDiet, Sandown Scientific, UK) that were automatically delivered and controlled by the VR software.

#### Behavioral paradigm

In our interval reproduction task, a rodent had to estimate the duration of a visual stimulus and reproduce it by moving along a virtual corridor. It is thus a variant of the “ready-set-go” timing task by Jazayeri and Shadlen (2010). Figure 1A illustrates the basic procedure: Each trial started with the presentation of a temporal stimulus – a black screen. Animals were trained to measure its duration and not to move during this phase of the task. Stimuli were randomly chosen between 3 and 7.5 seconds (i.e. either 3, 3.75, 4.5, 5.25, 6, 6.75, or 7.5 s). Afterwards, the visual scene switched, a virtual corridor appeared and the animal had to reproduce the stimulus by moving through the corridor for the same duration. The animal decided on its own when to start reproducing the interval as well as when to stop. These “reaction times” typically took a few seconds and only correlated weakly with the stimulus or the reproduced stimulus (Fig. S2C). If the animal continuously moved the treadmill for at least 1 s, the start of this movement was counted as the begin of reproduction. To finish reproduction, the animal had to stop for more than 0.5 s. These 0.5 s were not included in the reproduced interval. With this procedure, we avoided counting brief movements and stops as responses. Figure S1 shows movement data from an example session.

We gave feedback to our gerbils on their reproduction performance. Following the reproduction phase, the entire projection screen was either set to green (positive, “in” or “hit”) or white (negative, “out”) for 3-4 s. In addition, the animal was rewarded for a hit with one food pellet. For a hit, the reproduction had to be sufficiently close to the stimulus interval, i.e. (1 ± *k*) × stimulus. The width of this feedback range depended on the stimulus since errors increase with stimulus, i.e. scalar variability (cf. Jazayeri and Shadlen, 2010; Sohn et al., 2019). Across the session, tolerance *k* was reduced by −3% for hits and extended by +3% otherwise (Fig. 1B).

In the first trial of a session, *k* was always set to the value from the last trial in the previous session. Adapting *k* over a session, animals reached values close to 15% and below on average (Fig. 1E). Hit rates lay between 50% and 75% (Fig. S2B), indicating that the adaptive feedback range was successful in preventing alternative strategies such as learning the lower border of the feedback range that could also explain underestimation.

The virtual corridor was designed to exclude landmark-based strategies. It was infinite and had a width of 0.5 m. The walls of 0.5 m height were covered with a repetitive pattern of black and white stripes, each with a height to width ratio of 1:5. The floor was homogeneously colored in medium light-blue and the sky was black.

By randomly changing the gain between an animals’ own-movement (i.e. movement on the treadmill) and movement in VR, we de-correlated movement time from virtual distance and thus prevented path integration strategies for task solving. Gain values were uniformly sampled between 0.25 and 2.25. Distributions of virtual speed, running speed as well as their correlations with stimulus, reproduced stimulus and the bias (i.e. reproduction - stimulus) can be found in Fig. S2D&E. Running speed was (mostly negatively) correlated in about 25% of the sessions to stimulus and reproduction.

#### Behavioral training and testing

Naive gerbils were accustomed to the VR setup in a virtual linear maze for five to ten sessions (~ 2 weeks, cf. Thurley et al., 2014). Then, we exposed the animals to the timing task. As a first step, we presented only stimuli of 3 and 6 s which were easy to distinguish for the animals. The animals had to learn to either walk for a short or a long duration. Feedback was initially given with a tolerance of *k* = 50% and training proceeded until values below 30% were reached for at least three subsequent sessions. This training phase took about 1.5 months (about 30 sessions). In the second part of the training, we presented the full stimulus range for about 7 sessions (1.5 weeks), to introduce the animals to stimuli on a continuous scale. Afterwards, we implanted tetrodes into the animals’ mPFC and continued with the test phase.

#### Analysis of behavioral data

To compare behavioral performance across sessions and animals, we calculated different measures. To quantify the strength of the regression effect, we determined the slope of the linear regression between stimuli *s* and their reproductions *r*. A slope of one would correspond to no regression and smaller slopes to stronger regression. Variability is measured by the coefficient of variation, which we calculated as 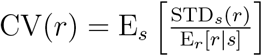. Here E_*r*_ [*r* | *s*] is the average response to a stimulus *s* and STD_*s*_(*r*) the corresponding standard deviation. The ratio of both values is averaged over all stimuli, denoted by E_*s*_[·]. To quantify general under- or overestimation, we use the BIAS(*r*) = E_*s*_[E_*r*_, [*r* | *s*] − *s*].

### Electrophysiological recordings

#### Electrode implantation

We chronically implanted gerbils with 8 tetrodes mounted to an microdrive that allowed for movement of all tetrodes together (Axona Ltd., St. Albans, UK). Tetrodes were made of 17 *μ*m platinum-iridium wires (California Fine Wire Co.). For surgery, we anesthetized an animal with an initial dose of medetomidine-midazolam-fentanyl (0.15 mg/kg, 7.5 mg/kg, 0.03 mg/kg, s.c.) and later maintained anesthesia by 2/3 doses every 2h. The animal was placed on a heating pad to keep body temperature at 37°C and fixated in a stereotactic unit (Stoelting Co.). After giving local analgesia of the skull with lidocaine (Xylocain, Astra Zeneca GmbH), we drilled a hole into the skull above the right mPFC and placed tetrodes at an initial depth of 700 *μ*m into the cortex (2.1 mm AP, −0.7 mm ML, −0.7 mm DV; Radtke-Schuller et al., 2016). To protect the exposed part of the brain, we used alginate (0.5% sodium alginate and 10% calcium chloride, Sigma-Aldrich) and paraffin wax. Further holes were drilled into the frontal, parietal and occipital bone to place small jewellers’ screws to help anchoring the microdrive to the skull with dental acrylic (iBond Etch, Heraeus Kulzer GmbH, Germany; Simplex Rapid, Kemdent, UK). One of the screws served as electrical ground. At the end of the surgery, anesthesia was antagonized with atipamezole-flumazenil-naloxone (0.4 mg/kg, 0.4 mg/kg, 0.5 mg/kg, s.c.). During surgery and for three postsurgical days, we gave meloxicam as a painkiller (0.2 mg/kg, s.c.). In addition, enrofloxacin antibiosis (Baytril, 10 mg/kg, s.c.) was done for five to seven postsurgical days. The animals were allowed to recover for at least three days after surgery before recordings started.

#### Recording procedures

Extracellular action potentials of single units were recorded at a rate of 32 kHz (Digital Lynx SX, Neuralynx, Inc.). Unit activity was band-pass filtered at 600 Hz to 6 kHz. Each tetrode could be recorded differentially, being referenced by one electrode of another tetrode or the ground connected to one of the jewellers’ screws. Recordings were done with Neuralynx’ data acquisition software Cheetah v5.6.3.

To sample of different neurons throughout the experimental period, we lowered the position of the tetrodes along the dorsoventral axis of the medial prefrontal cortex. Lowering was done for 50 *μ*m at the end of every second experimental session to allow for stabilization until the next experiment.

#### Reconstruction of tetrode placement

Tetrode placement was verified histologically postmortem. Animals received an overdose of sodium pentobarbital and were perfused intracardially with 4% paraformaldehyde. Brains were extracted and incubated in paraformaldehyde for 1-2 days. Afterwards, the brain was washed in 0.02 M phosphate buffered saline and coronal slices of 60-80 *μ*m thickness were obtained and stained either with Neutralred or DiI (D282), NeuroTrace 500/525 Green Fluorescent Nissl Stain, and DAPI – all stains from Thermo Fischer Scientific. Histology of all animals can be found in Figure S3A.

### Analysis of electrophysiological data

A total of 1842 mPFC neurons were recorded over 105 experimental sessions, each with on average more than 50 trials (Fig. S2A); animal 10526: 367 cells in 40 sessions; animal 11769: 677 cells in 32 sessions; and animal 11770: 798 cells in 33 sessions.

#### Spike sorting

Spike sorting was done offline in two steps. First, data was automatically clustered with KlustaKwik (v1.6). Afterwards, clusters were improved manually in 2D projections of waveform features incl. peak and valley, the difference between both, and energy, i.e. integral of the absolute value of the waveform, with MClust v4.3 (http://redishlab.neuroscience.umn.edu/MClust/MClust.html) under MATLAB2015b (The MathWorks, Inc.). See Figure S3 for example spike clusters.

Quality of spike sorting was assessed by calculating (1) rate of inter-spike-interval (ISI) violations (spikes with ISI < 1.5 ms are assumed to come from different neurons) and (2) calculating the fraction of spikes missing assuming a symmetric distribution of amplitudes (Hill, Mehta, and Kleinfeld, 2011, for details see). We excluded clusters with ISI violations > 0.5 and amplitude cutoff > 0.1. Both measures were calculated using the implementation in the Allen Institute ecephys spike sorting Python modules https://github.com/AllenInstitute/ecephys_spike_sorting. Also, only units with stable firing throughout a session entered further analysis. A unit was considered stable if spike counts in 1 min windows did not drop below four standard deviations from the mean session firing rate.

#### Spike density functions (SDFs)

We determined SDFs for each task phase separately. To calculate a SDF, spikes were either aligned at the begin or the end of the task phase. Then, spikes were counted in 100 ms windows for all trials at the same stimulus and divided by window width to gain firing rates. The windows were right aligned (looking into the past) to gain causal SDFs. To avoid edge effects due to response variability at the same stimulus, trials were scaled to the average response for the reproduction phase. During the measurement phase trials had the same duration by design. For visualization only (Figs. 2, S4-S6), SDFs were smoothed with a half Gaussian kernel of 3-bin standard deviation whose direction matched the right alignment of the window for spike counting, i.e. which was looking into the past.

For the analyses in Figures 3-7 and accompanying supplementary figures, SDFs were z-scored to account for cell-specific differences in firing rate. In addition, we resampled to same number of bins (time-normalized) the SDFs of different cells and for different stimuli, to be able to compare data from stimuli and responses of different duration.

For the population plots in Figure 3A-C, neurons were sorted by the angle between their demixed PCA scores for the first two time course-related PCs (see below for the description of the demixed PCA). This takes into account the full response profile instead of single features like the peak firing rate.

Control SDFs used in Figures 4 and S14 were generated by (1) shuffling SDFs for each stimulus across cells, i.e. “shuffled data”, and (2) shuffling single responses over time, i.e. “noise”.

#### Single cell and population correlations

In Figure 3 we report different Pearson correlations for the single cell and population data. All correlations were calculated on the time-normalized SDFs. Pairwise correlations were determined for the responses to all stimuli between all pairs of neurons (insets in Figure 3A-C). The population vector correlation in Fig. 3D was calculated between the activity of all neurons in one time bin during measurement and the corresponding time bin during reproduction, i.e. correlating columns in Figure 3A&C. In Figure 3E, single cell activity across all stimuli was correlated between measurement and reproduction, i.e. correlating rows in Figure 3A&C. In Figure 3F, population vector correlations were calculated for all stimulus pairs, i.e. correlating all points belonging to a particular stimulus in Fig. 3A or B to those belonging to another stimulus. Similarly, correlations for all pairs of stimuli were determined for each individual cell in Figure 3G.

#### Principal component analysis (PCA)

To gain a reduced representation of the collective population activity over time, we applied demixed and conventional PCA in the *T*-dimensional space where each dimension represents the firing rate at a different time point. A neuron’s activity pattern is hence represented as one single point in this space and a PC will be a component that has *T* points and evolves over time. The first PC will be the temporal pattern of activity that explains most variance; the second PC the one orthogonal to the first with second most variance and so on. By this method, PCs represent collective population activity over time. With demixed PCA, this population activity can be separated into components related to stimulus (stimulus time interval in our case) and those related to the overall time course of the population activity independent of the stimulus.

Demixed PCA was performed separately for measurement and reproduction on the SDFs of all recorded neurons aligned at the respective onsets (Fig. 5); see Kobak et al. (2016) for a detailed description. We used the demixed PCA implementation available at https://github.com/machenslab/dPCA. When we applied demixed PCA on data from individual animals results were similar (Figs. S8, S12, and S13). Conventional PCA was also done separately for measurement and reproduction and on the SDFs of all recorded neurons aligned at the respective onsets (Fig. S9).

Bootstrapping was done by performing demixed PCA on 1000 random subsets comprising each 10% of the whole data set, i.e. 184 neurons. Subsets were picked in a stratified way, i.e. accounting for the different numbers of cells recorded in each animal. The function StratifiedShuffleSplit from scikit-learn was used for picking the subsets. Results were similar for 5% and 20% subsets.

#### Tensor maximum entropy surrogates

To test whether population responses contained collective contributions beyond what is expected from pooling single neurons, we generated control surrogate data according to (Elsayed and Cunningham, 2017). Random tensor maximum entropy surrogate samples were drawn that preserved the stimulus tuning of single neurons, correlations of single cell firing rates across time and signal correlations across neurons. The implementation we used is available at https://github.com/gamaleldin/rand_tensor.

#### Categorization of response types

We categorized cells into different response types by their score values for specific principal components: time course-related PC 1 and stimulus PC 1 for measurement and time course-related PCs 1-3 and stimulus PC 1 for reproduction. Since the scores for those PCs had single peaked distributions, displaying no obvious clusters or response groups (Fig. S10), we used the following procedure to construct response categories: First, a cell’s responses in either task phase were reconstructed as a linear combination from the above mentioned PCs weighted by the respective PCA scores. Then the variance of the cell’s response was determined, which was explained by this reconstruction. If this explained variance was below the cumulative overall explained variance for the PCs (measurement: 6%, reproduction: 29%; cf. Fig. S8) the cell was assigned to the “unrelated activity” category; cf. Fig. S10. Otherwise, first the strongest PC (the one with the largest absolute scores) was found and then the angles between this PC and each of the other PCs were determined (calculated on the absolute values). If any of those angles was above 22.5°, the other component was counted as contributing (cf. Fig. S10C). In total, 2^2^ = 4 different response types were possible in the measurement phase and 2^4^ = 16 in the reproduction phase, i.e. categories ranging from “unrelated activity” with no overlap with any of the PCs to activity explained by all PCs used for categorization.

Categories were validated by categorizing every cell by its scores for each of the 1000 bootstrapped demixed PCAs described above. Finally, the category with maximum likelihood was assigned to the cell.

In addition, we compared the number of cells in each category to a random prediction. For that, we constructed random surrogate data by shuffling SDFs across stimuli and cells, performed demixed PCA on this data, categorized each “surrogate cell” and counted the number of cells in each category. From 1000 such shufflings we got distributions of by chance expected cell counts in each category, which we used to determine *p*-values for the count in the original data. A level of 5% was chosen and indicated as significant in Figure 6.

#### Time decoding

To decode elapsed time, we used multiple linear regression (Wiener filter; Glaser et al., 2017, https://github.com/KordingLab/Neural_Decoding) between time points and the spike responses (SDFs) of all neurons, i.e. the SDFs of each neuron were weighted such that elapsed time could be most precisely decoded. The SDFs of all neurons were combined in a matrix **R** with the individual SDFs as columns, such that the matrix had as many rows as time points and as many columns as neurons. Representing ongoing time as a vector **t**, the regression problem reads

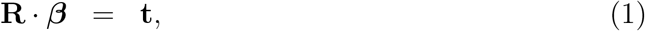

with *β* being the vector comprising the weights for each neuron (SDF), which can be fit by least squares.

The above equation deals only with one SDF per neuron, i.e. the response at one stimulus interval. However, we did not want to find individual weights for each stimulus but one weight for all stimuli per neuron. Therefore, we concatenated a neuron’s SDFs for all stimuli, such that **R** had as many rows as time points × stimuli. The weights *β* were then fit. This treated the whole data set as a reference (prior). During decoding ongoing time at a particular stimulus, we plugged in only the SDF at this stimulus into the left side of Eq. (1) and received a vector of decoded time points. Fitting and decoding was performed on random subsets of 20 cells (1000 bootstrap runs), from which we extracted average and standard deviation. To avoid overfitting, zero-mean Gaussian noise with *σ* = 0.5 was added to the SDFs. Note that the SDFs were z-scored.

### Additional notes on data analysis

Data analysis was done with Python 2.7 using Matplotlib 2.2, Numpy 1.15, Pandas 0.24, Scipy 1.2, Scikit-learn 0.20, and Statsmodels 0.10 – in addition to above mentioned packages. If p-values are not provided, significance is indicated by \**p* < 0.05; \*\**p* < 0.01; ***p <0.001.

## Acknowledgments

We thank Tobias Bernklau for help with data analysis and Moritz Dittmeyer for help with the schematic drawing of the setup. This work was supported by BMBF (Federal Ministry of Education and Research, Germany) via Bernstein Center Munich (grant number 01GQ1004A).

## Author contributions

KT and VFL envisioned the study and designed the behavioral paradigm. JH and KT designed and performed the experiments. KT, JH, DB, and DvW analyzed the data. KT and SH developed the method for categorization of response types. KT wrote the paper with help from the other authors.

## Declaration of Interests

The authors declare that they have no competing interests.

## Notes

### Competing Interest Statement

The authors have declared no competing interest.

