## Supplemental material for "Distributed coding of stimulus magnitude in rodent prefrontal cortex"

**References** **13**

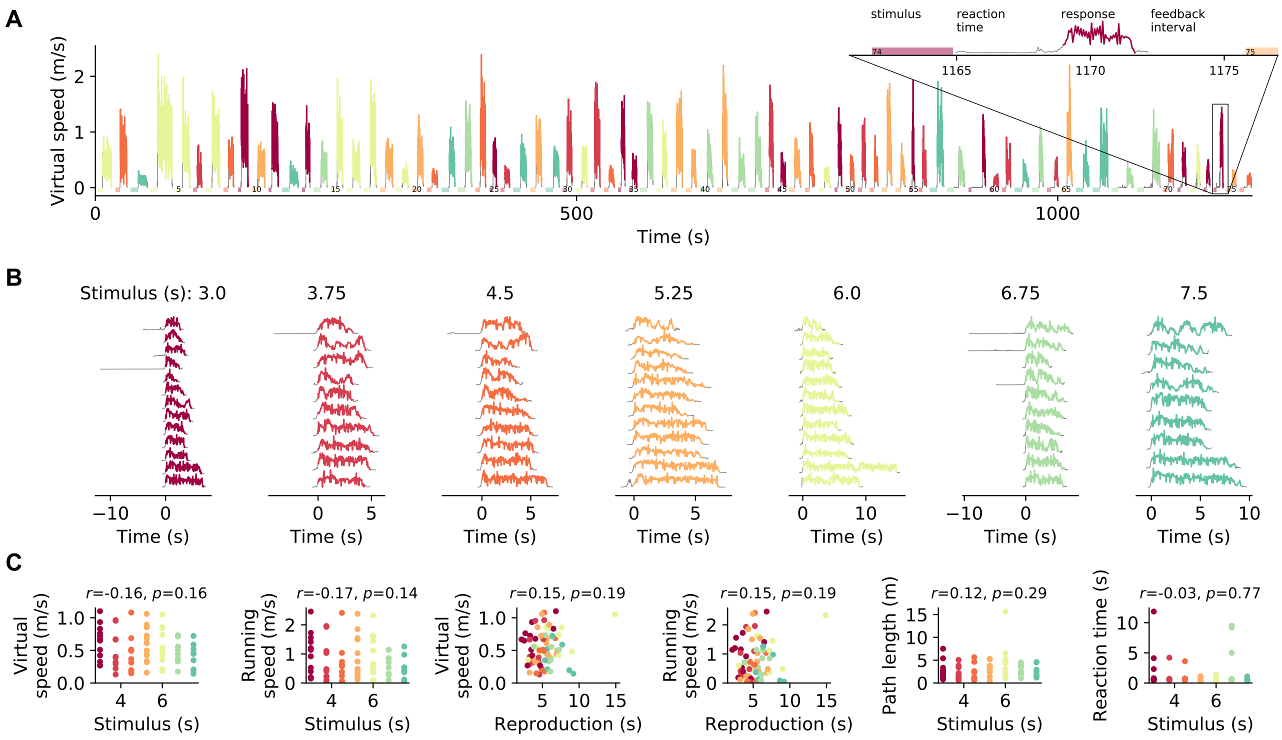

**Figure S1. Behavioral data from an example session.** (A) Virtual speed, i.e. the speed of visual movement an animal experiences, over the course of a session. Position was tracked during the response phase but not during measurement (stimulus) and feedback. Colored traces mark the intervals included as responses; gray otherwise. Color identifies stimulus. The interval between the end of stimulus presentation and the begin of the response we call "reaction time". To finish the response the animal had to stop for at least 0.5 s. *Inset:* Magnification of the 74th trial with labelled trial phases. (B) Virtual speed in individual trials sorted by stimulus. (C) Averages for each trial of the virtual speed, the running speed of the animal on the treadmill, the path length covered during the response, and the reaction time sorted by stimulus or reproduction. Again color identifies stimulus. Pearson's $r$ and $p$ values are given on top of each panel. The data belongs to the session displayed in the main text's Fig. 1D.

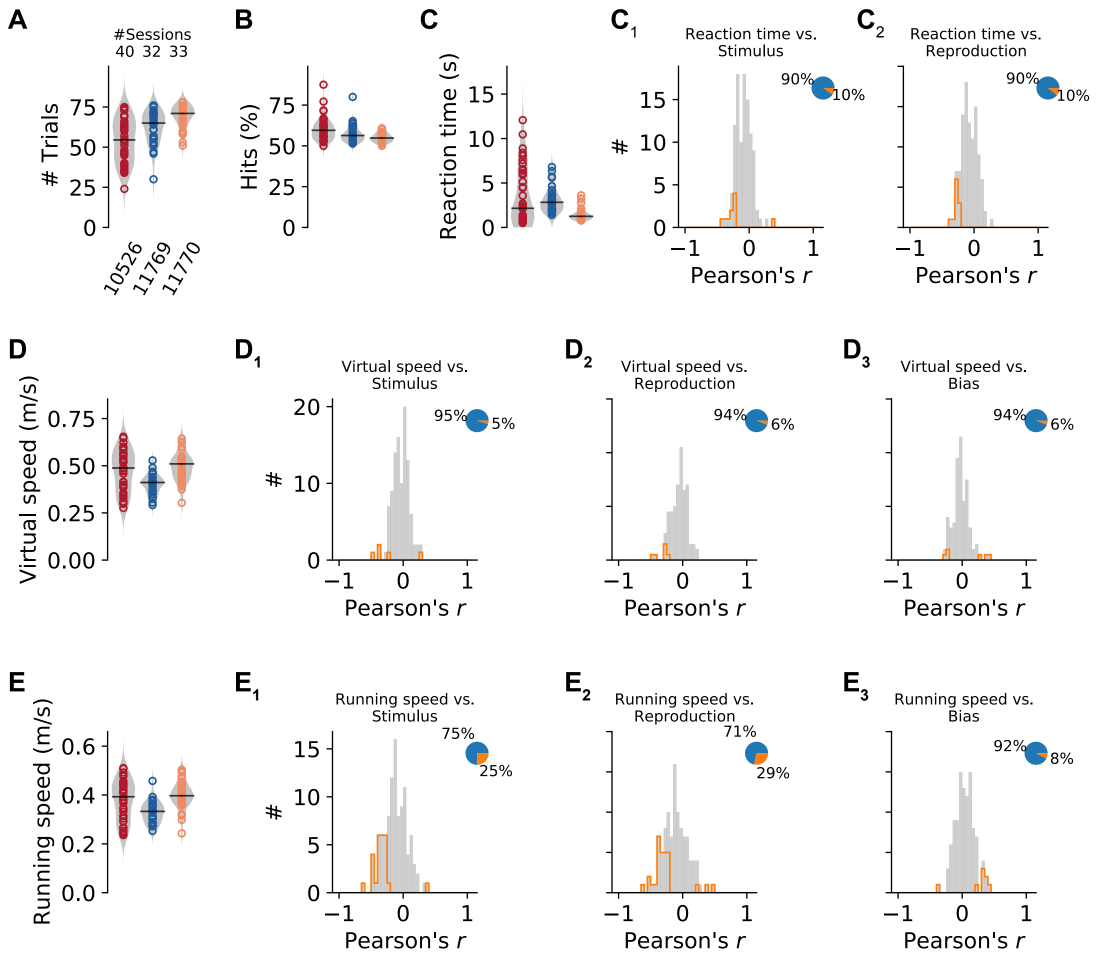

**Figure S2. Behavioral characteristics.** (A) Number of trials in the experimental sessions sorted by animal. Values from single sessions are displayed as open circles. Color identifies individual animals. Violin plots illustrate the distribution of the population. A solid black line marks the median. (B) Percentage of hits in the sessions. (C) Average "reaction time" in each session. ($C_1$&$C_2$) Distributions of Pearson's correlation coefficients between reaction time and stimulus or reproduced values. Histograms are displayed in gray with significant values delimited by an orange outline. Pie plots show significant (orange) and non-significant (blue) fractions. (D) Average virtual speed in each session and distributions of Pearson's correlations with stimulus ($D_1$), reproduction ($D_2$), and bias (i.e. reproduced value - stimulus, $D_3$). (E) Same as (D) for running speed of the animal on the treadmill.

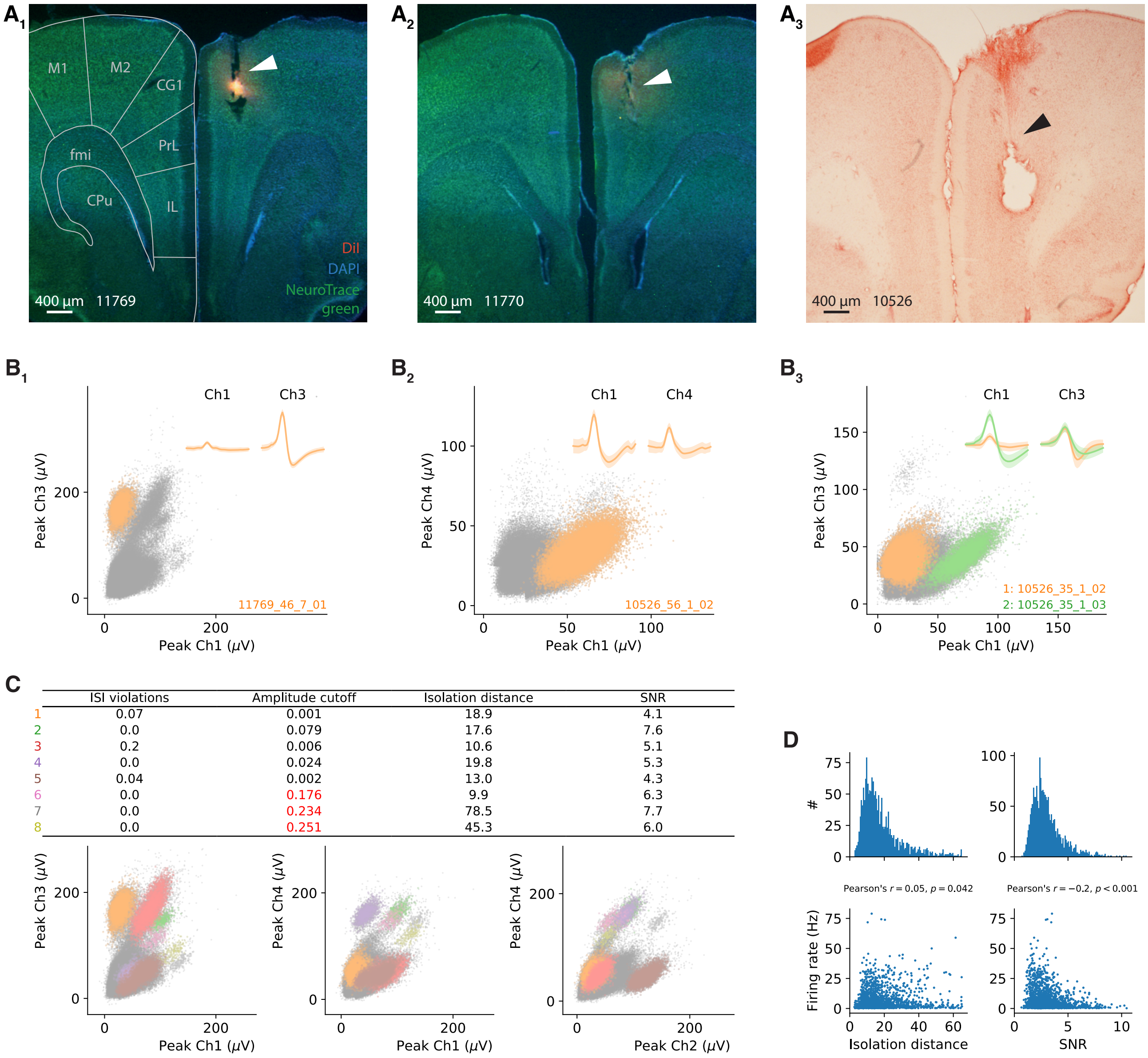

| | ISI violations | Amplitude cutoff | Isolation distance | SNR |
|---|---|---|---|---|
| 1 | 0.07 | 0.001 | 18.9 | 4.1 |
| 2 | 0.0 | 0.079 | 17.6 | 7.6 |
| 3 | 0.2 | 0.006 | 10.6 | 5.1 |
| 4 | 0.0 | 0.024 | 19.8 | 5.3 |
| 5 | 0.04 | 0.002 | 13.0 | 4.3 |
| 6 | 0.0 | 0.176 | 9.9 | 6.3 |
| 7 | 0.0 | 0.234 | 78.5 | 7.7 |
| 8 | 0.0 | 0.251 | 45.3 | 6.0 |

**Figure S3. Electrophysiological recordings in gerbil mPFC – histology and spike sorting.** (A) Representative coronal sections of the brains of all three gerbils. Tetrode tracks are visible in right cingulate cortex. In addition, stains from DiI coating of the tetrodes are visible in ($A_1$&$A_2$), and a big postmortem electrolytic lesion in ($A_3$). Background staining was done with DAPI and Neurotracer ($A_1$&$A_2$) and Neutralred ($A_3$). A section from a gerbil brain atlas is overlaid in ($a_1$) (Radtke-Schuller et al., 2016). M1, M2: primary and secondary motor cortex; Cg1: cingulate cortex, area 1; PrL: prelimbic cortex; IL: infralimbic cortex; fmi: forceps minor of corpus callosum; CPu: caudate putamen. (B) Spike clusters in feature space of mPFC cells recorded at one tetrode in three different sessions. ($B_1$) Cell from Fig. 2A in the main text. ($B_2$) cell from Fig. 2B in the main text. ($B_3$) cell from Fig. 2C (orange) in the main text and Fig. S5A (green). Peak signals are shown for two different channels of one tetrode as an example projection. Other spikes and voltage deflections are displayed as gray dots. *Insets:* Average waveforms (1 ms length) corresponding to the colored clusters. (C) Three different projections of the feature space from the tetrode recoding in ($B_1$). All eight spike clusters that could be separated are marked (color-coded). Other spikes and voltage deflections are again displayed as gray dots. The table lists quality measures for each cluster. Clusters 6 to 8 had too large amplitude cutoff and were excluded from further analysis. (D) Distributions of isolation distance and signal-to-noise ratio (SNR) for the clusters of all 1842 cells (i.e. excluding clusters/cells of insufficient quality). Both values were weakly correlated with the average firing rate (bottom panels). Isolation distance was calculated as described in Schmitzer-Torbert et al. (2005) for clusters of the peak amplitude (cf. (C)). SNR was calculated as the ratio of the peak of the average spike waveform to the standard deviation of the background noise, i.e. spikes that do not belong to a separated cluster. This was done for every channel individually and then averaged for all channels of a tetrode. ISI violations and amplitude cutoff were calculated as described in the Methods section.

### Example neurons

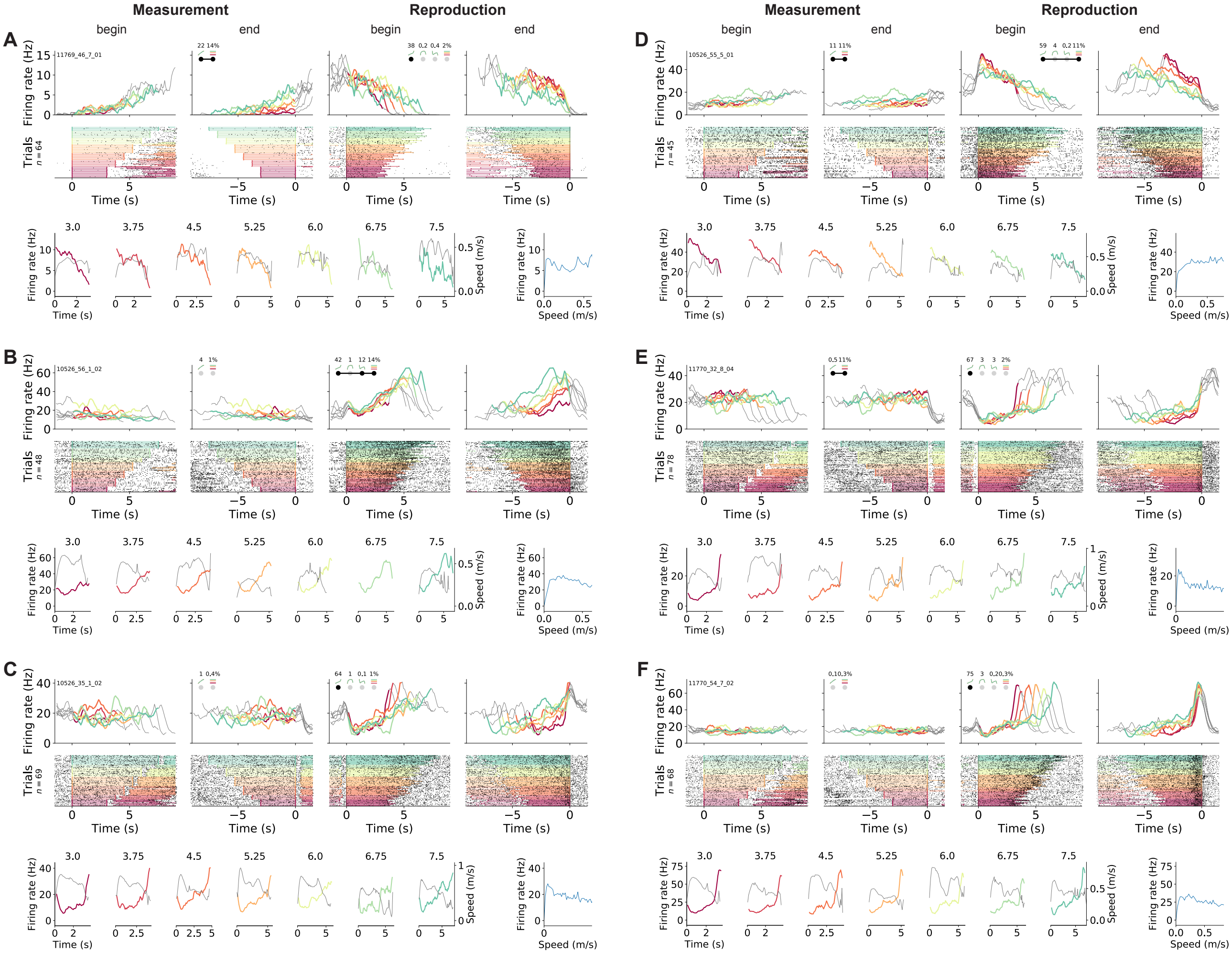

**Figure S4. Ramping neurons.** (A-C) Example cells from Fig. 2 in the main text. (D) Another cell that linearly increased its firing rate during measurement and ramped down to zero during reproduction. (E) A neuron that responded constantly but somewhat modulated by stimulus during measurement and ramps to threshold during reproduction. (F) Another ramp-to-threshold cell. (A-F) Panels display spike rasters sorted by stimulus (bottom) and corresponding spike density functions (SDF, top). Each column plots the data with different alignment, measurement begin and end, reproduction begin and end. Color identifies stimulus (cf. Fig. 1C). In the raster plots, black ticks are single spikes. For better visualization, we only plot half of the spikes (randomly chosen). Measurement or reproduction phases are delimited by underlayed color. The SDFs are colored in the respective task phase, outside they are displayed as thin gray lines. Markers in second and third panels show percent explained variance for each principal component (cf. Fig. 5). Black dots – that may be connected by a line – illustrate the cell type according to categorization (cf. Fig. 6). Bottom panels compare SDF (color-coded) to average (virtual) running speed (gray) for each stimulus. Bottom right panels plot the firing rate as a function of running speed. To calculate these speed response functions, we counted the number of spikes that occurred at a certain (virtual) running speed (2.5 cm/s bins) and divided the count by the total duration the animal moved at this speed. The response function is hence independent of trial and stimulus.

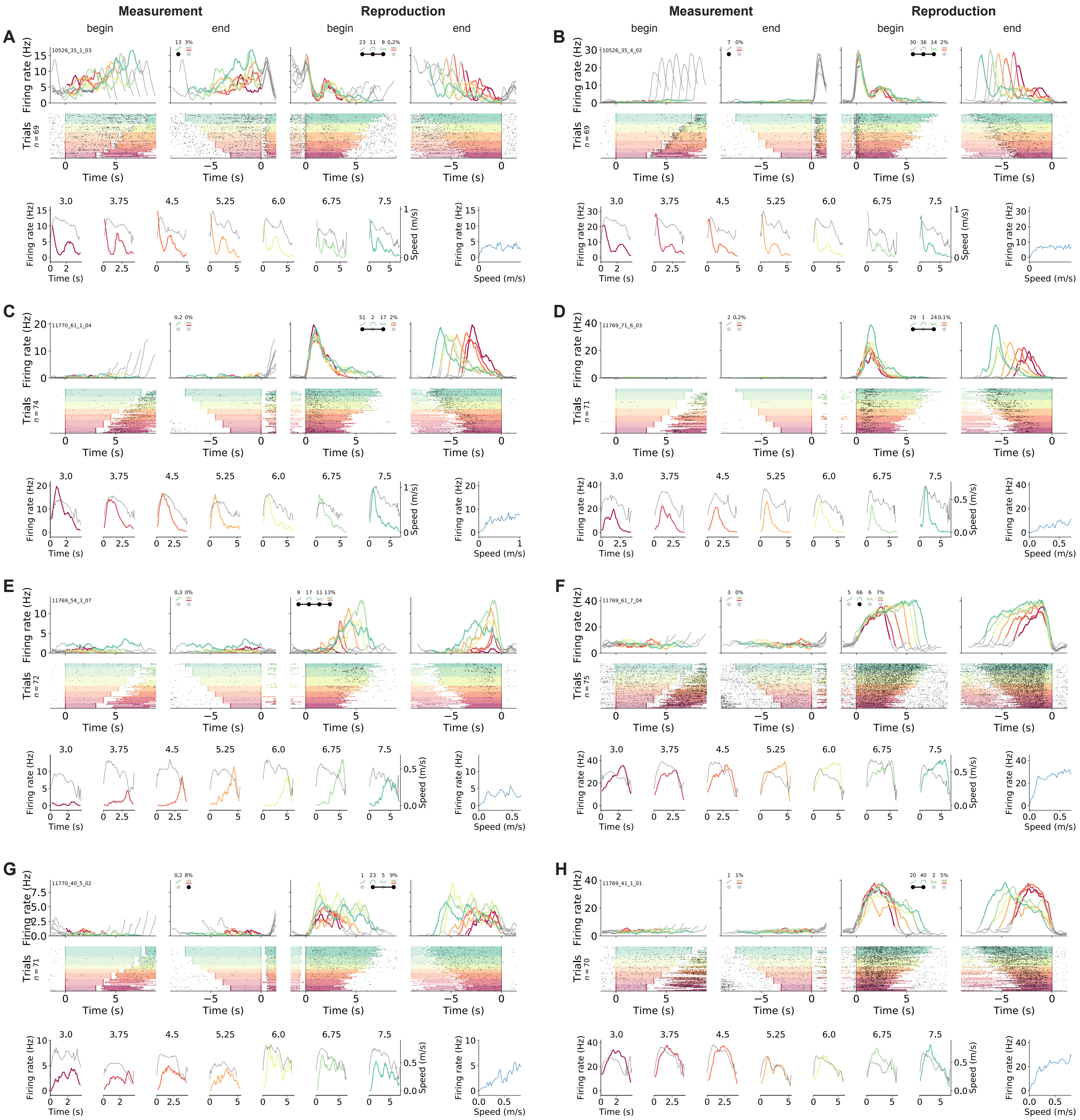

**Figure S5. Phasically timing neurons.** Examples of neurons that showed phasic responses at specific time points or for a certain duration in the reproduction phase. Plots are organized as in Fig. S4. Note, that the cell in (E) signals absolute time as it starts firing at  4 s – and proceeds until shortly before the end of the reproduced interval. (G&H) Neurons with pronounced modulation by (virtual) running speed. The shape of SDFs corresponds well to the profile of the running speed in these neurons. Also the neurons have a monotonously increasing speed response function. The reproduction SDFs of the neuron in (F) look very similar. However, its SDFs rather increase over the reproduction phase but the running speed decreases. This does not fit its speed response function. The response of the neuron therefore is determined by ongoing time rather than speed changes over a trial.

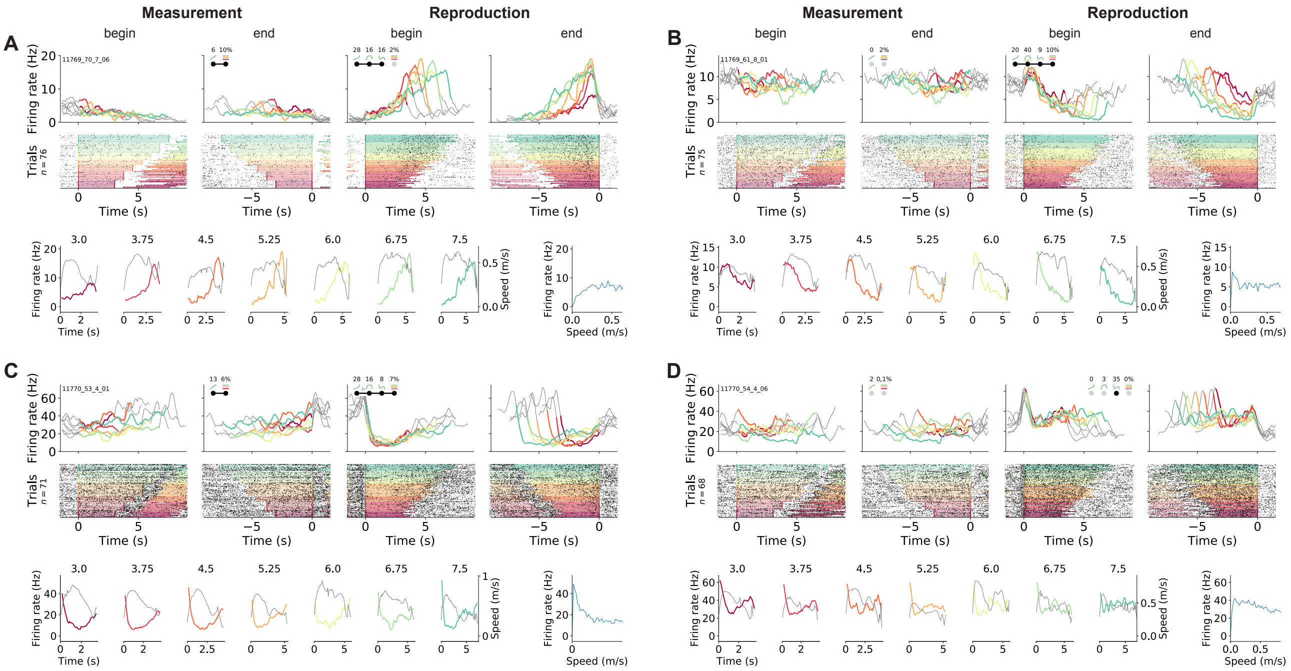

**Figure S6. Other example neurons.** (A) A neuron that during reproduction appeared very similar to the cell in Fig. S5F, but in fact does not signal absolute but relative time as it starts firing later for longer stimuli. Note the mild modulation by running speed, which saturates at ∼0.25 m/s, i.e. below the typical speed of the animal during the reproduction phase. (B-D) Cells that during reproduction responded with brief (D) or extended drops of activity (B&C). The amount of activity reduction may (C) or may not (B) have scaled with the stimulus. All plots are organized as in Fig. S4.

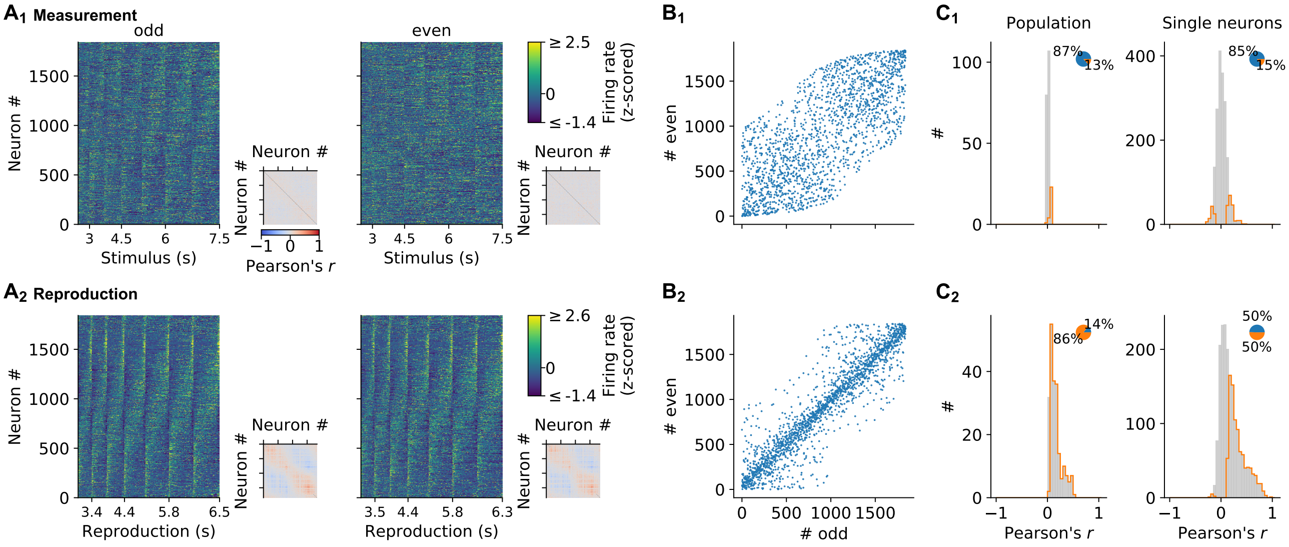

**Figure S7. Stability of single cell responses.** (A) Normalized (z-scored) SDFs of all neurons for each stimulus interval during ($A_1$) measurement and ($A_2$) reproduction sorted by their timing within the intervals; same sorting in ($A_1$) and ($A_2$). The left side displays data for odd trials only; the right panel for even trials. *Small panels:* Matrices of pairwise Pearson correlations between all neurons. Diagonal entries not plotted. (B) Indices assigned to the neurons if sorting would be based only on activity during odd or even trials. The closer a dot appears at the bisecting line, the better sorting corresponds between odd and even trials. During reproduction, activity corresponds well, arguing for stable responses throughout a recording session. During measurement activity is less similar. This has to be attributed to less pronounced and noisier responses in this task phase and not unstable recordings, since measurement and reproduction are interleaved throughout the session. (C) Distributions of Pearson correlations between odd and even trials give a similar picture. Histograms plot correlation coefficients during ($C_1$) measurement and ($C_2$) reproduction of the population in each time bin (left) and for single neurons (right). Histograms are displayed in gray with significant values delimited by an orange outline. Pie plots show significant (orange) and non-significant (blue) percentages.

### Supplementary figures related to principal component analysis

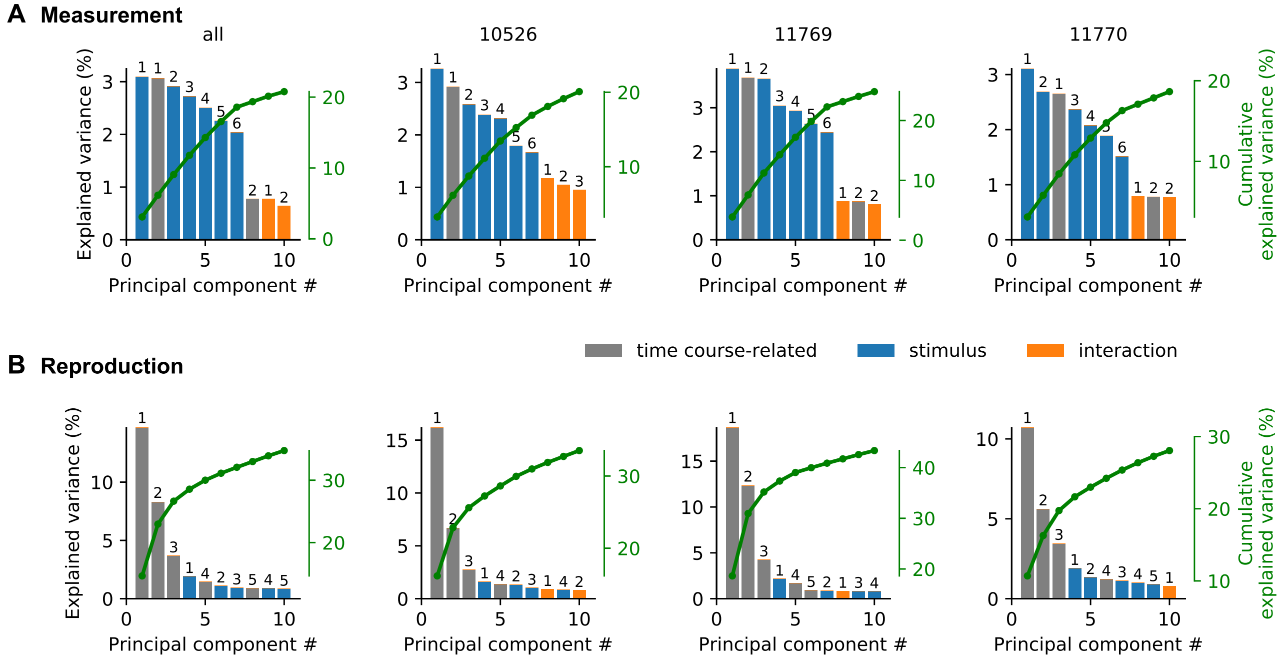

**Figure S8. Explained variance of demixed PCA.** Panels show explained variance for each principal component (bar graphs) and their cumulative explained variance (green line) of demixed PCA (Kobak et al., 2016) for measurement (A) and reproduction (B). Different panels correspond to all or individual animals. Numbers above bars give order with regard to component type, i.e. time course-related (gray), stimulus (blue), or interaction between both (orange). Results are comparable for individual animals and when pooled across all animals. In measurement the first time course-related and stimulus-dependent PCs are strongest. Reproduction is best explained by the first three time course-related PCs; only at forth place stimulus components start contributing.

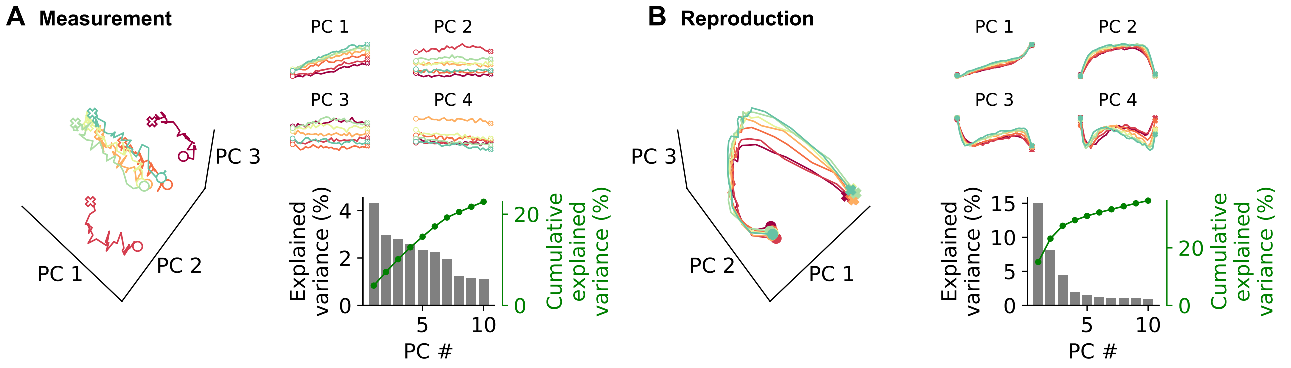

**Figure S9. Decomposition with conventional principal component analysis.** First four PCs for (A) measurement and (B) reproduction. Stimuli are colored as in other figures. Circles and crosses mark interval start and end. Open symbols are used for measurement and filled for reproduction. Bottom right panels: Explained variance for each principal component (bar graphs) and their cumulative explained variance (green line).

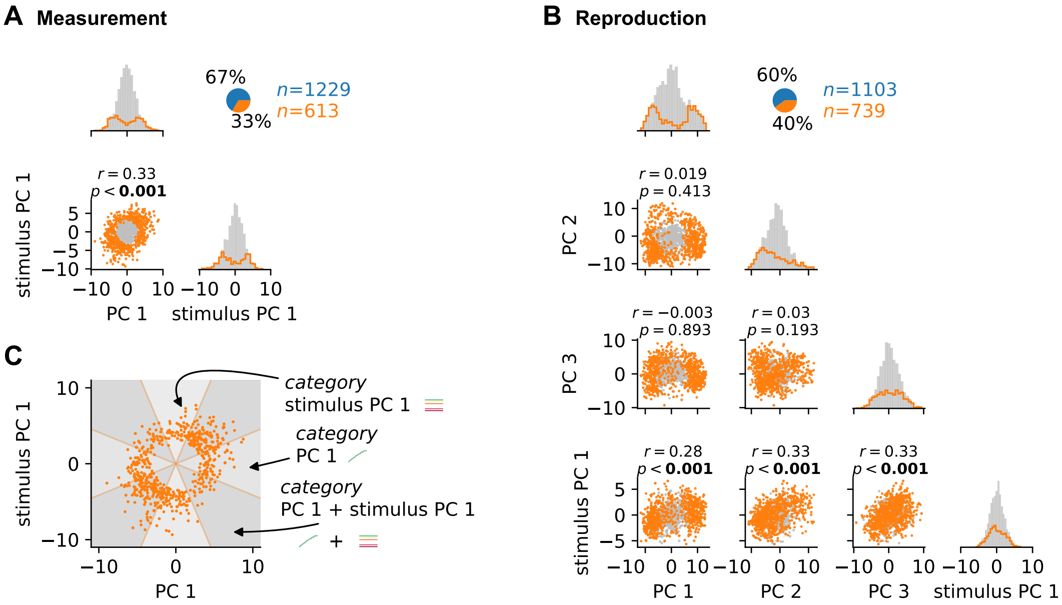

**Figure S10. Distributions of and correlations between demixed PCA scores.** (A) The scatter plot shows the correlation between scores for time course-related PC 1 and stimulus PC 1 in the measurement phase. Pearson's $r$ and $p$-value are listed above the scatter plot. Both were calculated for all cells. Histograms give distributions for the PCs. Cells that can be explained by the two PCs are plotted in orange, all others in gray. Pie plot shows fractions of cells that can (orange) and can not (blue) be explained by the principal components. (B) Same as (A) for reproduction and for time course-related PCs 1-3 and stimulus PC 1. (C) Categorization procedure at the example of the measurement data in (A). Categories were determined only for cells that could be sufficiently explained by the PCs. Cells where counted as explained by a PC if the ratio of scores (in absolute values) to the other category was below $\tan(67.5°)$. For the measurement phase, this results in three different categories: PC 1 only, stimulus PC 1 only, or PC 1 & stimulus PC 1 delineated by the three types of wedges in the plot.

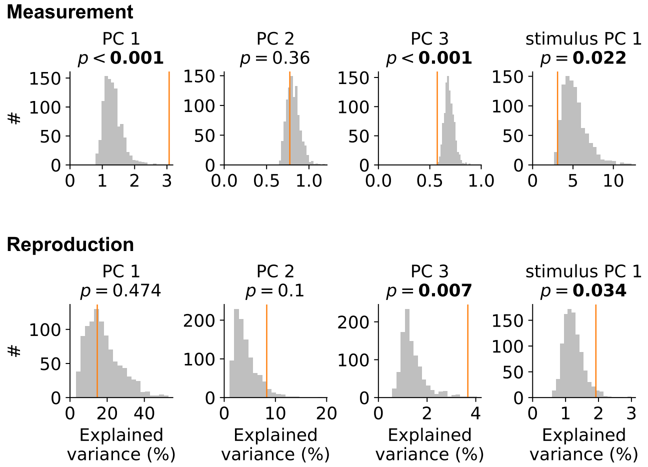

**Figure S11. Collective activity adds to PCs.** We drew 1000 random tensor maximum entropy surrogate samples (cf. Elsayed and Cunningham, 2017) for measurement (upper panels) and reproduction (lower panels) and determined the explained variance of time course-related components PC 1-3 and stimulus PC 1 (gray histograms). Orange vertical lines mark values for the real data. During measurement, PC 1 for the data was larger than expected. Similarly, PC 3 was larger than expected during reproduction. Stimulus PC 1 was larger than expectation for reproduction and at the lower end of what would be expected during measurement.

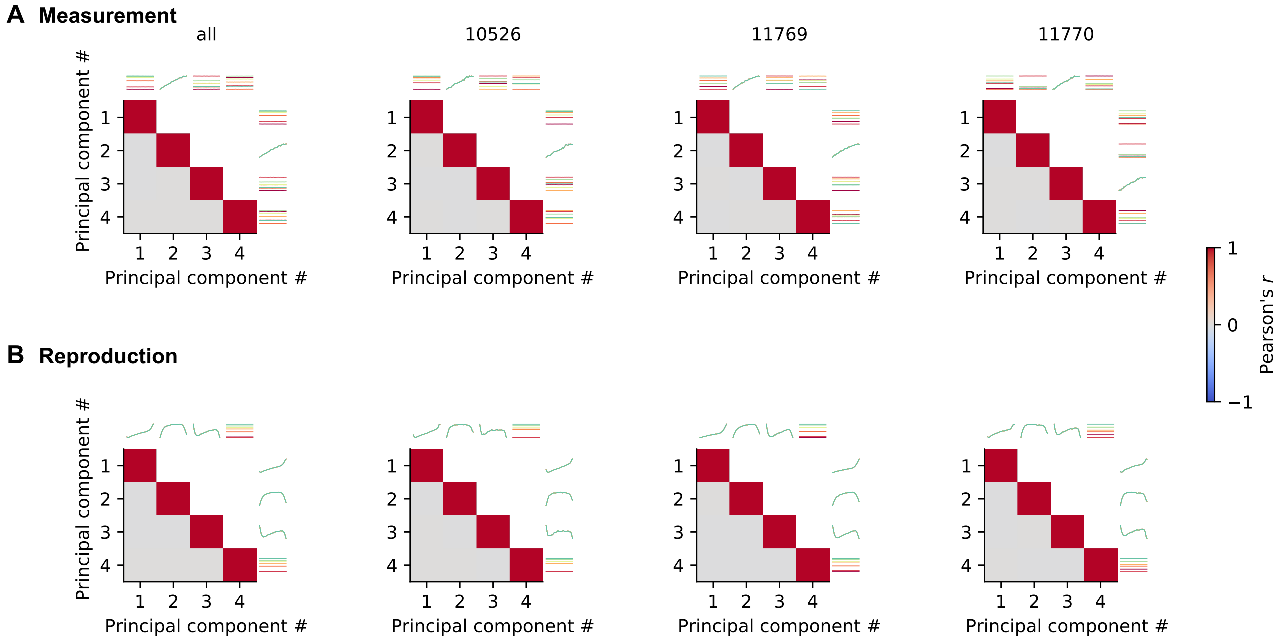

**Figure S12. Proper demixing of principal components by demixed PCA.** (A) Each panel shows correlations between all pairs of the first 4 demixed PCs for measurement. At top and to the left the principal components are illustrated. Different stimuli are color-coded as in other figures. Different panels give results for pooled data (all) and individual animals. (B) Same as (A) for reproduction. The correlations between different PCs are zero, indicating proper demixing.

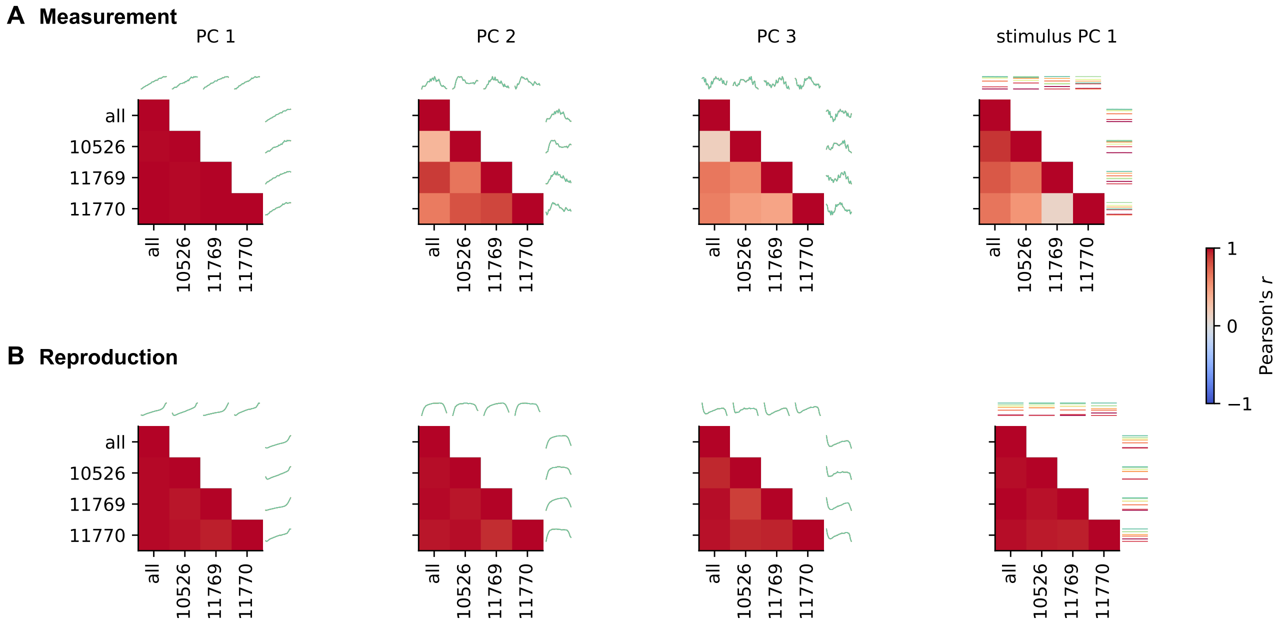

**Figure S13. Demixed PCA results are similar for each animal.** (A) Individual panels show correlations between all pairs of demixed PCs calculated for pooled data (all) and single animals for the measurement phase. At top and to the left the PCs are illustrated. Stimuli are color-coded. Different panels give time course-related PCs 1-3 and stimulus-related PC 1. (B) Same as (A) for reproduction. PCs are highly correlated across animals.

### Supplementary figures related to decoding

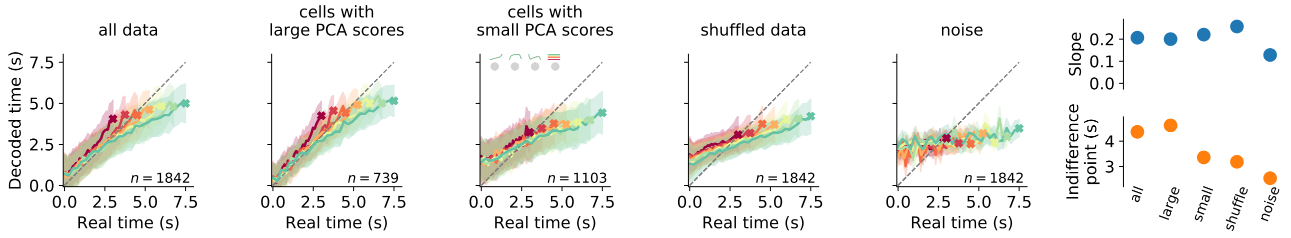

**Figure S14. Decoding elapsed time from data, shuffled data and noise.** Decoding results from the whole population, cells with sufficiently large dPCA scores, cells with small dPCA scores ("unrelated activity"), shuffled data and noise during the reproduction phase. Each panel displays decoded time vs. the real time for each stimulus (color-coded); average $\pm$ standard deviation (from bootstrapping). Crosses mark final values. Number of neurons are given in lower right corner. Rightmost panels display slopes and indifference points of linear regression between final values of real and predicted time for the five data sets. At the indifference point real and decoded final time match. We calculated it from the slope and intercept of the regression line as $\frac{\text{intercept}}{1-\text{slope}}$. The indifference point estimates the amount of general over-/underestimation. With a regression effect, the indifference point should be at the center of the stimulus distribution, which we only find when decoding from all data or the cells with large PCA scores.

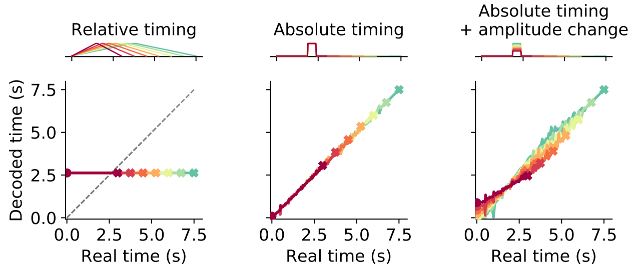

**Figure S15. Decoding time from phasically active neurons.** Decoding results for three different theoretical response types with phasic activation. The uppermost panel in each column shows example neuronal activity and the panel below gives the prediction for decoded time. Different stimulus intervals are color-coded. Crosses mark final values. Relative timing neurons peak at the center of an interval. Absolute timing neurons peak at a specific time point. In addition, stimulus may be coded in the response amplitude (Absolute timing + amplitude change). Although only one example is displayed, for both types of absolute timing neurons the population comprises cells peaking at different time points such that the whole interval is tiled. Relative timing neurons can only encode a single time point; whereas absolute timing neurons allow for precise time encoding.

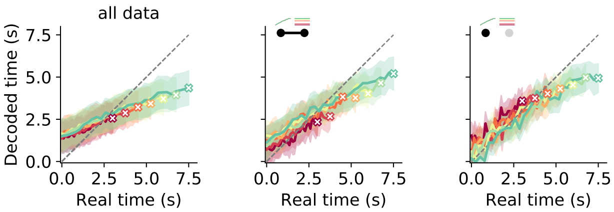

**Figure S16. Decoding time during measurement.** Results are displayed for all data (left), cells in the PC 1 + stimulus PC 1 category, and cells in the PC 1 only category. For all data, decoding is imprecise with initial overestimation and underestimation at the end of the interval. For PC 1 + stimulus PC 1-cells as well as cell in the PC 1 only-category decoded time starts out more precisely but ends-up at a general underestimation for the first response type. However, a regression effect with an overestimation for small stimuli is visible in the second case.
